## Supplementary Material 1 for "The value of livestock abortion surveillance in Tanzania: identifying disease priorities and informing interventions"

**SEBI Household Questionnaire [Dodoso la kaya]**

Enter household ID [Ingiza utambulisho wa kaya] [ ]
Language [Lugha]
☐ Kiswahili
☐ English
☐ Maa
Interviewer ID [Utambulisho wa mhojiwa] [ ]
Name of village in which household is located [Jina la kijiji] [ ]

**1. RESPONDENT/HOUSEHOLD HEAD DETAILS**
1.1. How many different households are in this compound [Kuna kaya ngapi katika boma hili]? [ ]

1.2 What is the relationship of the respondent to the head of the compound [Nini uhusiano wa mhojiwa na mkuu wa kaya]?
☐ Spouse [Mwenza]
☐ Parent [Mzazi]
☐ Child [Mtoto]
☐ Sibling [Dada/kaka]
☐ Self (= H of H) [Mimi mwenyewe]
☐ Other [Nyingine] [ ]

1.3 What is the head of compound's gender [Jinsia ya mkuu wa kaya]?
☐ Male [Mme] ☐ Female [Mke]

1.4 What is the head of compound's year of birth [Mwaka wako wa kuzaliwa ni upi]? [ ]

1.5 What is the head of compound's age class [Upo kwenye kundi gani la umri]?
☐ 13-18
☐ 19-34
☐ 35-54
☐ ≥55

1.6 What is the head of compound's tribe [Kabila lako ni lipi]?
☐ Arusha
☐ Barabaig
☐ Chagga
☐ Iraqw
☐ Pare
☐ Maasai
☐ Sambaa
☐ Other [ ]

1.7 What is the head of compound's highest level of education [Elimu yako cha juu ni ipi]?
☐ No formal education
☐ Some primary school
☐ Completed primary school
☐ Some secondary school
☐ Completed secondary school
☐ Post secondary qualifications
☐ Some university
☐ University completed
☐ Post graduate

**2. COMPOUND/HOUSEHOLD DETAILS**
2.1 How many members of this compound are in the following age-groups [Wakazi wangapi wa boma hili wako katika kila kundi la kiumri]?
< 5 years [chini miaka 5] [ ]
5-18 years [miaka 5-18] [ ]

> 18 years [zaidi ya miaka 18] [ ]

2.2 How many inhabited buildings are there in this compound [Kuna majengo mangapi ya kuishi katika kaya yako]? [ ]

2.3 What sources of drinking water does your compound use during the wet or dry season [Ni chanzo kipi cha maji kaya yako huwa wanatumia wakati wa masika]?
☐ Piped water into the home [Yanayosukumwa kwa bomba mpaka ndani nyumbani]
☐ Public/communal well or standpipe [Kisima au pampu ya jumuia]
☐ River or creek (moving water) directly [Moja kwa moja kutoka mto au mfereji (maji yanatotembea)]
☐ Lake, pond, dam (standing water) directly [Moja kwa moja kutoka Ziwa, dimbwi bwawa (maji yaliyosimama)]
☐ Private well or pump [Kisima au pampu ya binafsi]
☐ From a spring [Kutoka katika chemchem]
☐ Rainwater [Maji ya mvua]
☐ Tanker truck [Tanki la gari]
☐ Cart of wheelbarrow with small tank or drum [Mkokoteni na tanki dogo au madumu/pipa]
☐ Bottled water [Maji ya chupa]
☐ Other [ ] [Vyanzo vinginevyo]

2.4 How frequently do members of this compound treat water before drinking it [Huwa unatibu maji ya kunywa mara nyingi kabla ya kunywa]?
☐ Always (daily)
☐ Sometimes (a few times a week)
☐ Rarely (a few times a month)
☐ Never

2.5 How do you treat it [Ni mara ngapi huwa unatibu maji ya kunywa kabla ya kunywa]?
☐ Boiling [Kuchemsha]
☐ Strain it through a cloth [Kuchuja kwa nguo]
☐ Adding disinfectant, such as chlorine or bleach [Kuweka dawa kama shabu/klorine]
☐ Sedimentation and decant [Kuacha kwa muda yatwae/uchafu uende chini]

☐ Filtering [Kuchujwa]
☐ Solar disinfection [Kufanya salama kwa jua]
☐ Other [Nyinginezo] [ ]

2.6 What type of toilet system do members of this compound use [Ni aina gani ya mfumo wa choo ambao unatumika kawaida na kaya yako]?
☐ Flush or pour toilet with septic tank, including squat toilet [Choo cha kuvuta au cha kumwaga maji cha kuchuchumaa na mfumo wa shimo la maji taka]
☐ Flush or pour toilet connected to sewer pipe, including squat toilet [Choo cha maji kilichounganishwa na bomba la maji taka, pamoja na choo cha kuchuchumaa]
☐ Pit latrine with covering slab [Choo cha shimo kilichosakafiwa]
☐ Pit latrine without covering slab [Choo cha shimo bila kusakafiwa]
☐ Ventilated improved pit latrine (VIP) [Choo cha shimo bora chenya bomba la kutoa hewa chafu]
☐ Bucket or plastic bags [Ndoo au mifuko ya plastiki/Rambo]
☐ No facilities or field or bush [Hakuna choo, kwenda porini]
☐ Other [ ]

2.7 What kind of electricity do you have in this compound [Je, una umeme wa aina gani katika kaya hii]?
☐ Grid
☐ Solar
☐ Generator
☐ Other
☐ No electricity
☐ Other [ ]

2.8 What energy sources are used for cooking in this compound [Aina gani nishati inatumika kwa kupikia katika kaya hii]?
☐ Electricity [Umeme]
☐ Gas [Gesi]
☐ Kerosene [Kuni]
☐ Cow dung [Kinyesi cha ng'ombe]
☐ Firewood [Mafuta taa]
☐ Charcoal [Mkaa]
☐ Other [Nyinginezo] [ ]

2.9 For the house that the head of the compound sleeps in, what is the roof made of [Kwa nyumba anayolala mkuu wa kaya, paa limeezekwa na nini]?
☐ Metal [Chuma/mabati]
☐ Thatch [Nyasi]
☐ Wood [Mbao]
☐ Tiles [Vigae]
☐ Cement [Sementi]
☐ Dirt or mud [Vumbi/tope]
☐ Dung [Kinyesi cha mifugo]
☐ Other [Nyinginezo] [ ]

2.10 For the house that the head of the compound sleeps in, what is the floor made of [Kwa nyumba anayolala mkuu wa kaya, sakafu imetengenezwa na nini]?
☐ Brick [Matofali]
☐ Cement [Sementi]
☐ Wood [Mbao]
☐ Dirt or mud [Vumbi/tope]
☐ Dung [Kinyesi cha mifugo]
☐ Tile or linoleum [Vigae/sakafu ya mpira]
☐ Other [Nyinginezo] [ ]

2.11 For the house that the head of the compound sleeps in, what are the walls made of [Kwa nyumba anayolala mkuu kaya, ukuta umetengenezwa na nini]?
☐ Mud or manure [Tope au kinyesi cha mifugo]
☐ Burnt bricks [Tofali zilizochomwa]
☐ Mud bricks, uncooked [Tofali za matope]
☐ Cement bricks [Sementi]
☐ Wood [Mbao]
☐ Stone [Mawe]
☐ Thatch [Nyasi]
☐ Other [Nyinginezo] [ ]

2.12 Do members of this compound own any of the following items [Je, wakazi wa kaya hii wanamiliki chochote kati ya vitu vifuatavyo]?
☐ Ox plough [Chombo/kifaa]
☐ Ox cart [Mkokoteni wa ng'ombe]
☐ Bicycle [Baisikeli]
☐ Motorbike [Pikipiki]
☐ Car [Gari]
☐ Tractor [Trekta]
☐ Mobile phone [Simu ya mkononi]
☐ Radio [Redio]
☐ Television [Luninga]
☐ Sofa [Makochi]
☐ Refrigerator [Jokofu (Friji)]

**3. LAND AND CROPS**
3.1 Do members of this compound own land [Je, wanakaya wa kaya hii wanamliki ardhi]?
☐ Yes ☐ No

3.2 How much land do members of this compound own in total (in acres) [Ni kiasi gani cha ardhi wanakaya wa kaya hii wanamliki kwa ujumla (kwa eka)]? [ ]

3.3. Do members of the compound have a title for this land [Je, wanakaya wa kaya hii wana hati miliki ya hii ardhi]?
☐ Yes ☐ No

3.4 Where is the land title from [Je, hati miliki ya hii ardhi imetoka wapi]?
☐ Local (traditional)
☐ Government
☐ Other [ ]

3.5 Were any crops grown by members of this compound during the last 12 months [Kuna mazao yeyote yalioteshwa na watu wa kaya hili katika miezi 12 iliyopita]?
☐ Yes ☐ No

3.6 Which crops did members of the compound grow in the past 12 months [Kwa miezi 12 iliyopita ni mazao gani yalioteshwa na watu wa kaya hili]?
☐ Rice [Mpunga]
☐ Sorghum [Mtama]
☐ Sesame [Sesame]
☐ Sweet potato [Viazi vitamu]
☐ Cabbage [Kabeji]
☐ Tomato [Nyanya]
☐ Cotton [Pamba]
☐ Potato [Viazi]
☐ Spinach [Spinachi]
☐ Millet [Uwele]
☐ Maize [Mahindi]
☐ Cassava [Muhogo]
☐ Beans [Maharagwe]
☐ Lettuce [Saladi]
☐ Banana [Ndizi]
☐ Coffee [Kahawa]
☐ Avocado [Parachichi]
☐ Sugarcane [Miwa] ☐ Other [Nyinginezo]

3.7 How many years have members of this compound been growing crops [Ni kwa miaka mingapi watu wana kaya hii wameotesha mazao]? [ ]

3.8 Did this compound sell any crops in the last 12 months [Je, umeuza mazao yoyote katika miezi 12 iliyopita]?
☐ Yes ☐ No

**4. HOUSEHOLD LIVESTOCK OWNERSHIP**

4.1 Are any cattle currently kept at this compound [Je, kuna ng'ombe wowote wanafugwa kwenye boma hili kwa sasa]?
☐ Yes ☐ No

4.2 What is the total number of cattle that are managed together at this compound [Kuna jumla ya ng'ombe wangapi wanaotunzwa pamoja katika boma hili]? [ ]

4.3 Of these [NUMBER] cattle, how many are in the following breeds [Wangapi kati ya ngombe hawa]:
Indigenous [ ]
Crosses [ ]
Exotic [ ]

4.4 Of the [NUMBER] indigenous cattle, how many are the following types [Wangapi kati ya ngombe wakienyeji hawa]:
Adult male [ ]
Adult female [ ]
Juvenile male [ ]
Juvenile female [ ]

4.5 Of the [NUMBER] cross cattle, how many are the following types [Wangapi kati ya ngombe chotara hawa]:
Adult male [ ]

Adult female [ ]
Juvenile male [ ]
Juvenile female [ ]

4.6 Of the [NUMBER] exotic cattle, how many are the following types [Wangapi kati ya ngombe wa kisasa hawa]:
Adult male [ ]
Adult female [ ]
Juvenile male [ ]
Juvenile female [ ]

4.7 Are any goats currently kept at this compound [Je, kuna mbuzi wowote wanafugwa kwenye boma hili kwa sasa]?
☐ Yes ☐ No

4.8 What is the total number of goats that are managed together at this compound [Kuna jumla ya mbuzi wangapi wanaotunzwa pamoja katika boma hili]? [ ]

4.9 Of these [NUMBER] goats, how many are in the following breeds [Wangapi kati ya mbuzi hawa]:
Indigenous [ ]
Crosses [ ]
Exotic [ ]

4.10 Of the [NUMBER] indigenous goats, how many are the following types [Wangapi kati ya mbuzi wakienyeji hawa]:
Adult male [ ]
Adult female [ ]
Juvenile male [ ]
Juvenile female [ ]

4.11 Of the [NUMBER] cross goats, how many are the following types [Wangapi kati ya mbuzi chotara hawa]:
Adult male [ ]
Adult female [ ]
Juvenile male [ ]
Juvenile female [ ]

4.12 Of the [NUMBER] exotic goats, how many are the following types [Wangapi kati ya mbuzi wa kisasa hawa]:
Adult male [ ]
Adult female [ ]
Juvenile male [ ]
Juvenile female [ ]

4.13 Are any sheep currently kept at this compound [Je, kuna kondoo wowote wanafugwa kwenye boma hili kwa sasa]?
☐ Yes ☐ No

4.14 What is the total number of sheep that are managed together at this compound [Kuna jumla ya kondoo wangapi wanaotunzwa pamoja katika boma hili]? [ ]

4.15 Of these [NUMBER] sheep, how many are in the following breeds [Wangapi kati ya kondoo hawa]:
Indigenous [ ]
Crosses [ ]
Exotic [ ]

4.16 Of the [NUMBER] indigenous sheep, how many are the following types [Wangapi kati ya mbuzi wakienyeji hawa]:

Adult male [ ]
Adult female [ ]
Juvenile male [ ]
Juvenile female [ ]

4.17 Of these [NUMBER] cross sheep, how many are the following types [Wangapi kati ya kondoo chotara hawa]:
Adult male [ ]
Adult female [ ]
Juvenile male [ ]
Juvenile female [ ]

4.18 Of [NUMBER] exotic sheep, how many are the following types [ [Wangapi kati ya mbuzi wa kisasa hawa]::
Adult male [ ]
Adult female [ ]
Juvenile male [ ]
Juvenile female [ ]

**5. OTHER ANIMALS IN THE HOUSEHOLD**
5.1 Are any of the following animals owned in this compound [Kuna yoyote kati ya wanyama wafuatao wanamlikiwa katika boma hili]? [ ]
☐ Dogs [Mbwa]
How many dogs are there [Ni mbwa wangapi wapo]? [ ]
☐ Cats [Paka]?
How many cats are there [Ni paka wangapi wapo]? [ ]
☐ Donkeys [Punda]?
How many donkeys are there [Ni punda wangapi wapo]? [ ]
☐ Pigs [Nguruwe]?
How many pigs are there [Ni nguruwe wangapi wapo]? [ ]
☐ Chickens [Kuku]?
How many chickens are there [Ni kuku wangapi wapo]? [ ]
☐ Other birds [Ndege wengine]?
How many other birds are there [Ni ndege wengine wangapi wapo]? [ ]
☐ Other animals [Wanyama wengine]?
What are these other animals [Ni wanyama gani hao]? [ ]
How many [NUMBER] (other animals) are there [Hao [NUMBER](wanyama wengine) wako wangapi]? [ ]

**6. LIVESTOCK INTRODUCTIONS AND LOSSES**
6.1 How many cattle were born in this compound in the past 12 months [Ng'ombe wangapi wamezaliwa katika boma hili katika miezi 12 iliyopita]? [ ]

6.2 Have any cattle been introduced into this compound in the past 12 months? [Kuna ng'ombe yeyote aliyeliletwa katika kundi lenu kwa kipindi cha miezi 12 iliyopita]
☐ Yes ☐ No

6.3 How many of the introduced cattle were purchased [Wangapi kati yao walionunuliwa]? [ ]

6.4 How many of the introduced cattle were not purchased (e.g. a gift, dowry, exchange etc) [Wangapi hawakununuliwa au waliletwa kama zawadi, mahari, kubadalishana]? [ ]

6.5 Where did you buy the [NUMBER] cattle introduced into your household's herd [Kwa hao ng'ombe [NUMBER] walionunuliwa walitokea wapi]?
☐ Direct from another household
☐ Direct from market
☐ Livestock trader in this village (outside market)
☐ Livestock trader in another village (outside market)
☐ Don’t know
☐ Other [Nyinginezo] [ ]

6.6 Have any cattle kept in this compound died in the past 12 months (not through slaughter) [Kuna ng'ombe yoyote anayefugwa kwenye boma lako amekufa katika miezi 12 iliyopita (siyo kwa kuchinjwa)]?
☐ Yes ☐ No

6.7 What did these animals die of [Hawa wanyama walikufa na nini]?
Specify other [Vinginevyo (ainisha)]:
☐ Drought [Ukame]
☐ Predation [Kuliwa na mnyama]
☐ Disease [Ugonjwa]
☐ Trauma [Kuumia]
☐ Other [Nyinginezo] [ ]
☐ Don’t know [Sijui]

6.8 How many died from disease [Wangapi walikufa kutokana na magonjwa]? [ ]

6.9 How many goats were born in this household in the past 12 months [mbuzi wangapi wamezaliwa katika boma hili katika miezi 12 iliyopita]? [ ]

6.10 Have any goats been introduced into this household in the past 12 months? [Kuna mbuzi yeyote aliyeliletwa katika kundi lenu kwa kipindi cha miezi 12 iliyopita]
☐ Yes ☐ No

6.11 How many of the introduced goats were purchased [Wangapi kati yao walionunuliwa]?
[ ]

6.12 How many of the introduced goats were not purchased (e.g. a gift, dowry, exchange etc) [Wangapi hawakununuliwa au waliletwa kama zawadi, mahari, kubadalishana]? [ ]

6.13 Where did you buy the [NUMBER] goats introduced into your household's herd [Kwa hao mbuzi [NUMBER] walionunuliwa walitokea wapi]?
☐ Direct from another household
☐ Direct from market
☐ Livestock trader in this village (outside market)
☐ Livestock trader in another village (outside market)
☐ Don’t know
☐ Other [Nyinginezo] [ ]

6.14 Have any goats kept by your household died in the past 12 months (not through slaughter) [Kuna mbuzi yoyote anayefugwa kwenye boma lako amekufa katika miezi 12 iliyopita (siyo kwa kuchinjwa)]?
☐ Yes ☐ No

6.15 What did these animals die of [Hawa wanyama walikufa na nini]?
☐ Drought [Ukame]
☐ Predation [Kuliwa na mnyama]
☐ Disease [Ugonjwa]
☐ Trauma [Kuumia]
☐ Other [Nyinginezo] [ ]
☐ Don’t know [Sijui]

6.16 How many died from disease [Wangapi walikufa kutokana na magonjwa]? [ ]

6.17 How many sheep were born in this household in the past 12 months [kondoo wangapi wamezaliwa katika boma hili katika miezi 12 iliyopita]? [ ]

6.18 Have any sheep been introduced into this household in the past 12 months? [Kuna kondoo yeyote aliyeliletwa katika kundi lenu kwa kipindi cha miezi 12 iliyopita]
☐ Yes ☐ No

6.19 How many of the introduced sheep were purchased [Wangapi kati yao walionunuliwa]?
[ ]

6.20 How many of the introduced sheep were not purchased (e.g. a gift, dowry, exchange etc) [Wangapi hawakununuliwa au walileta kama zawadi, mahari, kubadalishana]? [ ]

6.21 Where did you buy the [NUMBER] sheep introduced into your household's herd [Kwa hao kondoo [NUMBER] walionunuliwa walitokea wapi]?
☐ Direct from another household
☐ Direct from market
☐ Livestock trader in this village (outside market)
☐ Livestock trader in another village (outside market)
☐ Don't know
☐ Other [ ]

6.22 Have any sheep kept by your household died in the past 12 months (not through slaughter) [Kuna kondoo yoyote anayefugwa kwenye boma lako amekufa katika miezi 12 iliyopita (siyo kwa kuchinjwa)]?
☐ Yes ☐ No

6.23 What did these animals die of [Hawa wanyama walikufa na nini]?
Specify other [Vinginevyo (ainisha)]:
☐ Drought [Ukame]
☐ Predation [Kuliwa na mnyama]
☐ Disease [Ugonjwa]
☐ Trauma [Kuumia]
☐ Other [Nyinginezo] [ ]
☐ Don't know [Sijui]

6.24 How many died from disease [Wangapi walikufa kutokana na magonjwa]? [ ]

**7. ANIMAL ILLNESS**
7.1 Have any animals in this compound been unwell (but not died) in the past 12 months?
☐ Yes ☐ No

Please tell us about the illness that affected the most cattle

7.2 How many cattle were affected [ ]

7.3 Which age group was affected?
☐ Only adults
☐ Only juveniles
☐ Adults and juveniles

7.4 Which sex was affected?
☐ Only females
☐ Only males
☐ Males and females

7.5 What signs did you observe?
☐ Milk drop
☐ Staring coat
☐ Regurgitation
☐ Diarrhoea
☐ Nasal discharge
☐ Coughing
☐ Listlessness
☐ Recumbancy
☐ Abdominal pain
☐ Jaundice
☐ Inappetance
☐ Lameness
☐ Weight loss
☐ Other [ ]

7.6 Do you know what the cause was?

☐ Yes ☐ No

7.7 What was the cause? [ ]
7.8 How many months ago did this occur? [ ]

7.9 Were there any other (different) episodes of illness in cattle?
☐ Yes ☐ No

7.10 How many cattle were affected? [ ]

7.11 Which age group was affected?
☐ Only adults
☐ Only juveniles
☐ Adults and juveniles

7.12 Which sex was affected?
☐ Only females
☐ Only males
☐ Males and females

7.13 What signs did you observe?
☐ Milk drop
☐ Staring coat
☐ Regurgitation
☐ Diarrhoea
☐ Nasal discharge
☐ Coughing
☐ Listlessness
☐ Recumbancy
☐ Abdominal pain
☐ Jaundice
☐ Inappetance
☐ Lameness
☐ Weight loss
☐ Other [ ]

7.14 Do you know what the cause was?
☐ Yes ☐ No

7.15 What was the cause? [ ]

7.16 How many months ago did this occur? [ ]

7.17 Were there any other (different) episodes of illness in cattle?
☐ Yes ☐ No

7.18 How many cattle were affected? [ ]

7.19 Which age group was affected?
☐ Only adults
☐ Only juveniles
☐ Adults and juveniles

7.20 Which sex was affected?
☐ Only females
☐ Only males
☐ Males and females

7.21 What signs did you observe?
☐ Milk drop
☐ Staring coat
☐ Regurgitation
☐ Diarrhoea
☐ Nasal discharge
☐ Coughing
☐ Listlessness
☐ Recumbancy
☐ Abdominal pain
☐ Jaundice
☐ Inappetance
☐ Lameness
☐ Weight loss
☐ Other [ ]

7.22 Do you know what the cause was?
☐ Yes ☐ No

7.23 What was the cause? [ ]

7.24 How many months ago did this occur? [ ]

Please tell us about the illness that affected the most goats

7.25 How many goats were affected? [ ]

7.26 Which age group was affected?
☐ Only adults
☐ Only juveniles
☐ Adults and juveniles

7.27 Which sex was affected?
☐ Only females
☐ Only males
☐ Males and females

7.28 What signs did you observe?
☐ Milk drop
☐ Staring coat
☐ Regurgitation
☐ Diarrhoea
☐ Nasal discharge
☐ Coughing
☐ Listlessness
☐ Recumbancy
☐ Abdominal pain
☐ Jaundice
☐ Inappetance
☐ Lameness
☐ Weight loss
☐ Other [ ]

7.29 Do you know what the cause was?
☐ Yes ☐ No

7.30 What was the cause? [ ]

7.31 How many months ago did this occur? [ ]

7.32 Were there any other (different) episodes of illness in goats?
☐ Yes ☐ No

7.33 How many goats were affected? [ ]

7.34 Which age group was affected?
☐ Only adults
☐ Only juveniles
☐ Adults and juveniles

7.35 Which sex was affected?
☐ Only females
☐ Only males
☐ Males and females

7.36 What signs did you observe?
☐ Milk drop
☐ Staring coat
☐ Regurgitation
☐ Diarrhoea
☐ Nasal discharge
☐ Coughing
☐ Listlessness
☐ Recumbancy
☐ Abdominal pain
☐ Jaundice
☐ Inappetance
☐ Lameness
☐ Weight loss
☐ Other [ ]

7.37 Do you know what the cause was?
☐ Yes ☐ No

7.38 What was the cause? [ ]

7.39 How many months ago did this occur? [ ]

7.40 Were there any other (different) episodes of illness in goats?
☐ Yes ☐ No

7.41 How many goats were affected? [ ]

7.42 Which age group was affected?
☐ Only adults
☐ Only juveniles
☐ Adults and juveniles

7.43 Which sex was affected?
☐ Only females
☐ Only males
☐ Males and females

7.44 What signs did you observe?
☐ Milk drop
☐ Staring coat
☐ Regurgitation
☐ Diarrhoea
☐ Nasal discharge
☐ Coughing
☐ Listlessness
☐ Recumbancy
☐ Abdominal pain
☐ Jaundice
☐ Inappetance
☐ Lameness
☐ Weight loss
☐ Other [ ]

7.45 Do you know what the cause was?
☐ Yes ☐ No

7.46 What was the cause? [ ]

7.47 How many months ago did this occur? [ ]

Please tell us about the illness that affected the most sheep

7.48 How many sheep were affected? [ ]

7.49 Which age group was affected?
☐ Only adults
☐ Only juveniles
☐ Adults and juveniles

7.50 Which sex was affected?
☐ Only females
☐ Only males
☐ Males and females

7.51 What signs did you observe?
☐ Milk drop
☐ Staring coat
☐ Regurgitation
☐ Diarrhoea
☐ Nasal discharge
☐ Coughing
☐ Listlessness
☐ Recumbancy
☐ Abdominal pain
☐ Jaundice
☐ Inappetance
☐ Lameness

☐ Weight loss
☐ Other [ ]

7.52 Do you know what the cause was?
☐ Yes ☐ No
7.53 What was the cause? [ ]

7.54 How many months ago did this occur? [ ]

7.55 Were there any other (different) episodes of illness in sheep?
☐ Yes ☐ No

7.56 How many sheep were affected? [ ]

7.57 Which age group was affected?
☐ Only adults
☐ Only juveniles
☐ Adults and juveniles

7.58 Which sex was affected?
☐ Only females
☐ Only males
☐ Males and females

7.59 What signs did you observe?
☐ Milk drop
☐ Staring coat
☐ Regurgitation
☐ Diarrhoea
☐ Nasal discharge
☐ Coughing
☐ Listlessness
☐ Recumbancy
☐ Abdominal pain
☐ Jaundice
☐ Inappetance
☐ Lameness
☐ Weight loss
☐ Other [ ]

7.60 Do you know what the cause was?
☐ Yes ☐ No

7.61 What was the cause? [ ]

7.62 How many months ago did this occur? [ ]

7.63 Were there any other (different) episodes of illness in sheep?
☐ Yes ☐ No

7.64 How many sheep were affected? [ ]

7.65 Which age group was affected?
☐ Only adults
☐ Only juveniles
☐ Adults and juveniles

7.66 Which sex was affected?
☐ Only females
☐ Only males
☐ Males and females

7.67 What signs did you observe?
☐ Milk drop
☐ Staring coat
☐ Regurgitation
☐ Diarrhoea
☐ Nasal discharge
☐ Coughing
☐ Listlessness
☐ Recumbancy
☐ Abdominal pain
☐ Jaundice

☐ Inappetance
☐ Lameness
☐ Weight loss
☐ Other [ ]

7.68 Do you know what the cause was?
☐ Yes ☐ No

7.69 What was the cause? [ ]

7.70 How many months ago did this occur? [ ]

7.71 Has milk production changed in any species in the past 12 months?
☐ Cattle [Ng'ombe]
☐ Goats [Mbuzi]
☐ Sheep [Kondoo]

7.72 How has it changed in cattle?
☐ Increased
☐ Decreased
☐ Don't know

7.73 How has it changed in goats?
☐ Increased
☐ Decreased
☐ Don't know

7.74 How has it changed in sheep?
☐ Increased
☐ Decreased
☐ Don't know

7.75 Have any species had joint problems in the past 12 months?
☐ Yes ☐ No

**8. LIVESTOCK MANAGEMENT**

8.1 Have any animals in your herd been vaccinated in the past 24 months [Je, kuna wanyama katika kundi wamepata chanjo katika kipindi cha miezi 24 iliyopita]?
☐ Yes ☐ No

8.2. Against which diseases [Ni chanjo dhidi ya magonjwa gani]?
☐ CBPP
☐ Anthrax
☐ FMD
☐ LSD
☐ RVF
☐ Sheep pox
☐ Goat pox
☐ Other [ ]

8.3 How are cattle in this compound grazed during the wet and dry season [Unawalishaje ng'ombe wa boma hili kipindi cha mvua na kiangazi]?
Free ranging [Wanajichunga wenyewe]: ☐ Dry ☐ Wet ☐ Neither
Herded [Wanachungwa]: ☐ Dry ☐ Wet ☐ Neither
Tethered [Wanaofungwa]: ☐ Dry ☐ Wet ☐ Neither
Zero grazed [Hawachungwi]: ☐ Dry ☐ Wet ☐ Neither

8.4 How are goats in this compound grazed during the wet and dry season [Unawalishaje mbuzi wa boma hili kipindi cha mvua na kiangazi]?
Free ranging [Wanajichunga wenyewe]: ☐ Dry ☐ Wet ☐ Neither
Herded [Wanachungwa]: ☐ Dry ☐ Wet ☐ Neither

Tethered [Wanaofungwa]: ☐ Dry ☐ Wet ☐ Neither
Zero grazed [Hawachungwi]: ☐ Dry ☐ Wet ☐ Neither

8.5 How are sheep in this compound grazed during the wet and dry season [Unawalishaje kondoo wa boma hili kipindi cha mvua na kiangazi]?

Free ranging [Wanajichunga wenyewe]: ☐ Dry ☐ Wet ☐ Neither
Herded [Wanachungwa]: ☐ Dry ☐ Wet ☐ Neither
Tethered [Wanaofungwa]: ☐ Dry ☐ Wet ☐ Neither
Zero grazed [Hawachungwi]: ☐ Dry ☐ Wet ☐ Neither

8.6 Which of the following best describes the way you manage the herding of animals in this compound [Ipi kati yafuatayo inaeleza vizuri jinsi unavyotunza kundi la wanyama katika boma hili]:

☐ Cattle, sheep and goats together [BO, OV, na CP pamoja]
☐ Cattle separately, sheep and goats together [OV na CP pamoja]
☐ Cattle with goats, sheep separately [BO na CP pamoja]
☐ Cattle with sheep, goats separately [BO na OV pamoja]
☐ All species separately [Kila aina inachungwa tofauti]
☐ Other [Nyinginezo] [ ]

8.7 How long do your livestock walk (in minutes) from this compound for grazing on a normal day during the dry season? [Inachukuwa muda gani wa mifugo kutembea kwa ajili ya malisho (kwa dakika) kutoka boma hili wakati wa kiangazi]

Cattle [Ng'ombe] [ ]
Goats [Mbuzi] [ ]
Sheep [Kondoo] [ ]

8.9 How long do your livestock walk (in minutes) from this compound for grazing on a normal day during the wet season [[Inachukuwa muda gani wa mifugo kutembea kwa ajili ya malisho (kwa dakika) kutoka boma hili wakati wa mvua]?

Cattle [Ng'ombe] [ ]
Goats [Mbuzi] [ ]
Sheep [Kondoo] [ ]

8.10 How long do your livestock walk (in minutes) from this compound for water on a normal day during the dry season [Inachukuwa muda gani wa mifugo kutembea kwa ajili ya maji (kwa dakika) kutoka boma hili wakati wa kiangazi]?

Cattle [Ng'ombe] [ ]
Goats [Mbuzi] [ ]
Sheep [Kondoo] [ ]

8.11 How long do your livestock walk (in minutes) from this compound for water on a normal day during the wet season [Inachukuwa muda gani wa mifugo kutembea kwa ajili ya maji (kwa dakika) kutoka boma hili wakati wa masika]?

Cattle [Ng'ombe] [ ]
Goats [Mbuzi] [ ]
Sheep [Kondoo] [ ]

8.12 Are cattle from this compound regularly taken to seasonal camps for grazing [Je, ng'ombe wa boma hili wanapelekwa kwenye maboma ya muda kwa ajili ya malisho (ronjo)]?

☐ Yes ☐ No

8.13 Where are these seasonal camps (for cattle) [Haya maboma ya muda yapo wapi (kwa ng'ombe)]?

Village or place name [Kijiji au jina la sehemu] [ ]
Ward [Kata] [ ]
District [Wilaya] [ ]

8.14 Out of the [NUMBER] cattle in this compound, how many would normally go to seasonal grazing [Kati ya ng'ombe [NUMBER] ambao wako katika boma hii ni wangapi kwa kawaida wanaenda kwenye maboma ya muda kwa malisho]? [ ]

8.15 Are goats from this compound regularly taken to seasonal camps for grazing [Je, mbuzi wa boma hili wanapelekwa kwenye maboma ya muda kwa ajili ya malisho (ronjo)]?
☐ Yes ☐ No
8.16 Is this to the same place as for cattle [Sehemu hili ni sawa sawa za sehemu za n'gombe]?
☐ Yes ☐ No

8.17 Where are these seasonal camps (for goats)?
Village or place name [Kijiji au jina la sehemu] [ ]
Ward [Kata] [ ]
District [Wilaya] [ ]

8.18 Out of the [NUMBER] goats in this compound, how many would normally go to seasonal grazing [Kati ya mbuzi [NUMBER] ambao wako katika boma hli ni wangapi kwa kawaida wanaenda kwenye maboma ya muda kwa malisho]? [ ]

8.19 Are sheep from this compound regularly taken to seasonal camps for grazing [Je, kondoo wa boma hili wanapelekwa kwenye maboma ya muda kwa ajili ya malisho (ronjo)]?
☐ Yes ☐ No

8.20 Are these the same place as for cattle and goats [Sehemu hili ni sawa sawa za sehemu za n'gombe na mbuzi]?
☐ Yes ☐ No

8.21 Are these the same place as for cattle [Sehemu hili ni sawa sawa za sehemu za n'gombe]?
☐ Yes ☐ No

8.22 Are these the same place as for goats [Sehemu hili ni sawa sawa za sehemu za mbuzi]?
☐ Yes ☐ No

8.23 Where are these seasonal camps (for sheep) [Haya maboma ya muda yapo wapi (kwa kondoo)]?
Village or place name [Kijiji au jina la sehemu] [ ]
Ward [Kata] [ ]
District [Wilaya] [ ]

8.24 Out of the [NUMBER] sheep in this compound, how many would normally go to seasonal grazing [Kati ya kondoo [NUMBER] ambao wako katika boma hii ni wangapi kwa kawaida wanaenda kwenye maboma ya muda kwa malisho]? [ ]

8.25 Which of the following best describes the way you manage animals at night in this household [Ipi kati ya yafutatayo inaeleza vizuri jinsi unavyotunza wanyama wako katika kaya hii wakati wa usiku]:
☐ Cattle, sheep and goats together [BO, OV, na CP pamoja]
☐ Cattle separately, sheep and goats together [OV na CP pamoja]
☐ Cattle with goats, sheep separately [BO na CP pamoja]
☐ Cattle with sheep, goats separately [BO na OV pamoja]
☐ All species separately [Kila aina inachungwa tofauti]
☐ Other [Nyinginezo] [ ]

**9. BREEDING, PARTURITION, AND ABORTION**
9.1 How was service of livestock done in this compound in the past 12 months [Huduma ya upandishaji wa mifugo ilitolewa namna gani katika kundi lako kwa miezi 12 uliopita]?

| | | | |
|---|---|---|---|
| Males from own herd: [Madume (dume) kutoka kundi lako] | ☐ Cattle | ☐ Goats | ☐ Sheep |
| Males hired: [Madume (dume) wakukodisha] | ☐ Cattle | ☐ Goats | ☐ Sheep |
| Males borrowed: [Madume (dume) ya kuazima] | ☐ Cattle | ☐ Goats | ☐ Sheep |
| Males from other herds during grazing/watering: [Wanaume kutoka wanyama wa wengine / kumwagilia] wakati wa kulima | ☐ Cattle | ☐ Goats | ☐ Sheep |
| Artificial insemination: [Uzalishaji kwa chupa] | ☐ Cattle | ☐ Goats | ☐ Sheep |
| Females taken elsewhere: [Jike lilipelekwa pengine] | ☐ Cattle | ☐ Goats | ☐ Sheep |
| No service performed in past 12 months: [Hakuna ushirikiano uliofanywa katika miezi 12 iliyopita] | ☐ Cattle | ☐ Goats | ☐ Sheep |
| Other [Nyinginezo]: | ☐ Cattle | ☐ Goats | ☐ Sheep |
| Species not owned or service not performed: | ☐ Cattle | ☐ Goats | ☐ Sheep |

Specify other for cattle [ ]
Specify other for goats [ ]
Specify other for sheep [ ]

9.2 In the past 12 months, where were the young livestock owned by this compound born? [Je, indani wa miezi 12 iliyopita, mifugo wanao milikiwa na boma hili wamezaliwa wapi]?

| | | | |
|---|---|---|---|
| In a (human) house in this compound : | ☐ Cattle | ☐ Goats | ☐ Sheep |
| In the compound (outside a house) : | ☐ Cattle | ☐ Goats | ☐ Sheep |
| Outside the compound in this village : | ☐ Cattle | ☐ Goats | ☐ Sheep |
| Outside the compound in a different village : | ☐ Cattle | ☐ Goats | ☐ Sheep |
| Other: | ☐ Cattle | ☐ Goats | ☐ Sheep |
| Species not owned/no births in past 12 months: | ☐ Cattle | ☐ Goats | ☐ Sheep |

Specify other for cattle [ ]
Specify other for goats [ ]
Specify other for sheep [ ]

9.3 Do cattle that have just given birth usually get seperated from the main herd [Je, n'gombe ambao wamezaa wanatenganisha na kundi/wanyama wengine]?
☐ Yes ☐ No

9.4 For how long [Kwa muda gani]?
☐ < 24 hours
☐ 12 to 24 hours
☐ > 24 hours

9.5 How many days [Kwa siku ngapi]? [ ]

9.6 Do goats that have just given birth usually get seperated from the main herd [Je, mbuzi ambao wamezaa wanatenganisha na kundi/wanyama wengine]?
☐ Yes ☐ No

9.7 For how long [Kwa muda gani]?
☐ < 24 hours
☐ 12 to 24 hours
☐ > 24 hours

9.8 How many days [Kwa siku ngapi]? [ ]

9.9 Do sheep that have just given birth usually get seperated from the main herd [Je, kondoo ambao wamezaaa wanatenganisha na kundi/wanyama wengine]?
☐ Yes ☐ No

9.10 For how long [Kwa muda gani]?
☐ < 24 hours
☐ 12 to 24 hours
☐ > 24 hours

9.11 How many days [Kwa siku ngapi]? [ ]

9.12 What normally happens to the placenta and foetal membranes after one of your animals (sheep, cattle, or goats) gives birth [Kwa kawaida inakuwaje kwa kondo la nyuma baada ya mojawapo ya wanyama wako (ng'ombe, mbuzi, au kondoo) wanapozaa]?
☐ Nothing, material left [Hakuna – viliachwa]
☐ Eaten by family [Vililiwa na wakazi katika familia]
☐ Buried [Vilifukiwa (zikwa)]
☐ Burned [Vilichomwa]
☐ Given to dogs [Vilipewa mbwa vikiwa vibichi
☐ Thrown in bush [Vilitupwa porini]
☐ Other [Nyinginezo] [ ]

9.13 How many cattle in this compound have aborted or delivered still born offspring in the past 12 months [Ni ng'ombe wangapi katika kaya yako wametupa mimba au kuzaa mtoto mfu katika miezi 12 iliyopita]? [ ]

9.14 How many goats in this compound have aborted or delivered still born offspring in the past 12 months [Ni mbuzi wangapi katika kaya yako wametupa mimba au kuzaa mtoto mfu katika miezi 12 iliyopita]? [ ]

9.15 How many sheep in this compound have aborted or delivered still born offspring in the past 12 months [Ni kondoo wangapi katika kaya yako wametupa mimba au kuzaa mtoto mfu katika miezi 12 iliyopita]? [ ]

9.16 What normally happens to the foetus, placenta and/or membranes after one of your livestock has an abortion or stillbirth [Kwa kawaida inakuwaje kwa kichanga, kondo la nyuma baada ya moja ya mifugo wako kutupa mimba au kuzaa mtoto mfu]?
☐ Nothing, material left [Hakuna – viliachwa]
☐ Eaten by family [Vililiwa na wakazi katika familia]
☐ Buried [Vilifukiwa (zikwa)]
☐ Burned [Vilichomwa]
☐ Given to dogs [Vilipewa mbwa vikiwa vibichi
☐ Thrown in bush [Vilitupwa porini]
☐ Other [Nyinginezo] [ ]

9.17 What do you normally do with an animal after it has an abortion [Kwa kawaida unachukuwa hatua gani kwa mnyama baada ya kutupa mimba]?

| | | | |
|---|---|---|---|
| Nothing : | ☐ Cattle | ☐ Goats | ☐ Sheep |
| Separate animal from herd for <24 hours : | ☐ Cattle | ☐ Goats | ☐ Sheep |
| Separate animal from herd for >24 hours : | ☐ Cattle | ☐ Goats | ☐ Sheep |
| Send animal for slaughter : | ☐ Cattle | ☐ Goats | ☐ Sheep |
| Sell animal : | ☐ Cattle | ☐ Goats | ☐ Sheep |
| Give animal away : | ☐ Cattle | ☐ Goats | ☐ Sheep |
| Get advice/treatment from outside household : | ☐ Cattle | ☐ Goats | ☐ Sheep |
| Treat the animal myself : | ☐ Cattle | ☐ Goats | ☐ Sheep |
| Other : | ☐ Cattle | ☐ Goats | ☐ Sheep |
| Never owned or had an abortion in this species : | ☐ Cattle | ☐ Goats | ☐ Sheep |

Specify other for cattle [ ]
Specify other for goats [ ]
Specify other for sheep [ ]

9.18 Do you think you would be more likely to sell animals that have had an abortion than those that have not [Je, unafikiri kuna uwezekano mkubwa kuuza wanyama aliyetupa mimba kuliko wale ambao hawakutupa mimba]?

Cattle [Ng'ombe] [ ]
Goats [Mbuzi] [ ]
Sheep [Kondoo] [ ]

9.19 Do you think you would be more likely to slaughter animals that have had an abortion than those that have not [Je, unafikiri kuna uwezekano mkubwa kuchinja wanyama aliyetupa mimba kuliko wale ambao hawakutupa mimba]?
Cattle [Ng'ombe] [ ]
Goats [Mbuzi] [ ]
Sheep [Kondoo] [ ]

9.20 What treatment do you give when an animal has an abortion [Ni tiba gani gani unampa mnyama baada ya kutupa mimba]? [ ]

9.21 Who do you normally go to for advice/treament when an animal has an abortion [Ni nani kwa kawaida unakwenda kupata ushauri/matibabu kuhusu mimba ikitoka]?
☐ Neighbour [Jirani]
☐ Paravet [Paravet]
☐ Veterinarian [Bwana mifugo]
☐ Animal health assistant [Bwana afya msaidizi wa mifugo]
☐ Agrovet [Bwana kilimo na mifugo]
☐ Traditional healer [Mganga wa jadi]
☐ Spiritual/faith healer [Imani/mponyaji wa imani]
☐ Local livestock expert [Mtaalamu wa mifugo wa hapahapa]
☐ Other [Nyinginezo] [ ]

9.22 Have you ever had a case of retained fetal membranes in livestock kept in this compound [Je, kulishawai kuwa na matukio ya kondo la nyuma kuacha kutoka katika mifugo inayotunzwa katika boma hili (ikihusisha kaya yako)]?
☐ Yes ☐ No

9.23 How many cattle in this compound have had a case of retained fetal membrane in the past 12 months [Ni ng'ombe wangapi wa kundi la kaya yako wamepata matukio ya kondo la nyuma kuacha kutoka katika miezi 12 iliyopita]? [ ]

9.24 How many goats in this compound have had a case of retained fetal membrane in the past 12 months [Ni miezi mingapi iliyopita mara ya mwisho kuwepo tukio la kondo la nyuma kuacha kutoka katika kundi la mbuzi wa kaya hii]? [ ]

9.25 How many sheep in this compund have had a case of retained fetal membrane in the past 12 months [Ni kondoo wangapi wa kundi la kaya yako wamepata matukio ya kondo la nyuma kuacha kutoka katika miezi 12 iliyopita]? [ ]

9.26 What actions do you take when an animal has a retained placenta [Je, unachukua hatua gani kama mnyama akishindwa kutoa kondo la nyuma]?

| | | | |
|---|---|---|---|
| Nothing: | ☐ Cattle | ☐ Goats | ☐ Sheep |
| Separate animal from herd for <24 hours: | ☐ Cattle | ☐ Goats | ☐ Sheep |
| Separate animal from herd for >24 hours: | ☐ Cattle | ☐ Goats | ☐ Sheep |
| Send animal for slaughter: | ☐ Cattle | ☐ Goats | ☐ Sheep |
| Sell animal: | ☐ Cattle | ☐ Goats | ☐ Sheep |
| Give animal away: | ☐ Cattle | ☐ Goats | ☐ Sheep |
| Get advice/treatment from outside household: | ☐ Cattle | ☐ Goats | ☐ Sheep |
| Treat the animal myself: | ☐ Cattle | ☐ Goats | ☐ Sheep |
| Never owned/had retained fetal membrane: | ☐ Cattle | ☐ Goats | ☐ Sheep |
| Other [Nyinginezo] [ ] | | | |

9.27 Do you think you would be more likely to sell animals that have had a case of retained placenta than those that have not [Je, unafikiri kuna uwezekano mkubwa kuuza wanyama akishindwa kutoa kondo la nyuma kuliko wale ambao wametoa kondo la nyuma]?
Cattle [Ng'ombe] [ ]

Goats [Mbuzi] [ ]
Sheep [Kondoo] [ ]

9.28 Do you think you would be more likely to slaughter animals that have had a case of retained placenta than those that have not [Je, unafikiri kuna uwezekano mkubwa kuchinja wanyama akishindwa kutoa kondo la nyuma kuliko wale ambao wametoa kondo la nyuma]?
Cattle [Ng'ombe] [ ]
Goats [Mbuzi] [ ]
Sheep [Kondoo] [ ]

9.29 What treatment do you give when an animal in your herd has a retained placenta [Ni tiba gani unampa mnyama aliyeshindwa kutoa kondo la nyuma katika kundi lako]? [ ]

9.30 Who do you normally go to for advice/treament when your animal has a retained placenta [Kwa kawaida unakwenda kwa nani kupata ushauri/matibabu wakati mnyama wako ameshindwa kutoa kondo la nyuma]?
☐ Neighbour [Jirani]
☐ Paravet [Paravet]
☐ Veterinarian [Bwana mifugo]
☐ Animal health assistant [Bwana afya msaidizi wa mifugo]
☐ Agrovet [Bwana kilimo na mifugo]
☐ Traditional healer [Mganga wa jadi]
☐ Spiritual/faith healer [Imani/mponyaji wa imani]
☐ Local livestock expert [Mtaalamu wa mifugo wa hapahapa]
☐ Other [Nyinginezo] [ ]

9.31 Have you identified infertile male or female livestock in this compound in the past 12 months [Je, umegunduwa mfugo wowote dume au jike amabye ni tasa katika boma hili miezi 12 iliyopita (ikihusisha boma lako)]?
☐ Yes ☐ No

9.32 How many male cattle have been identified as infertile in the past 12 months [Ni madume mangapi ya ng'ombe wa kundi la kaya yako wamegundulika kuwa tasa miezi 12 iliyopita]? [ ]

9.33 How many female cattle have been identified as infertile in the past 12 months [Ni majike mangapi ya ng'ombe wa kundi la kaya yako wamegundulika kuwa tasa miezi 12 iliyopita]? [ ]

9.34 How many male goats have been identified as infertile in the past 12 months [Ni madume mangapi ya mbuzi wa kundi la kaya yako wamegundulika kuwa tasa miezi 12 iliyopita]? [ ]

9.35 How many female goats have been identified as infertile in the past 12 months [Ni majike mangapi ya mbuzi wa kundi la kaya yako wamegundulika kuwa tasa miezi 12 iliyopita]? [ ]

9.36 How many male sheep have been identified as infertile in the past 12 months [Ni madume mangapi ya kondoo wa kundi la kaya yako wamegundulika kuwa tasa miezi 12 iliyopita]? [ ]

9.37 How many female have been identified as infertile in the past 12 months [Ni majike mangapi ya kondoo wa kundi la kaya yako wamegundulika kuwa tasa miezi 12 iliyopita]? [ ]

9.38 What actions do you take when you discover an animal is infertile [Unachukua hatua gani kwa kawaida wakati mnyama ni tasa]?
Nothing : ☐ Cattle ☐ Goats ☐ Sheep
Kinachofanyika-wanyama walioadhirika wanatunzwa kama wengine
Send animal for slaughter: ☐ Cattle ☐ Goats ☐ Sheep

[Kuchinja]
Sell animal: ☐ Cattle ☐ Goats ☐ Sheep
[Kuuza]
Give animal away: ☐ Cattle ☐ Goats ☐ Sheep
Kupeanwa
Get advice/treatment from outside household: ☐ Cattle ☐ Goats ☐ Sheep
Ushauri au matibabu yanatafutwa nje ya kaya
Treat the animal myself:
[Kutibu mnyama mwenyewe] ☐ Cattle ☐ Goats ☐ SheepOthe
Never owned or had infertility in this species:
[Haijawahi kumiliki au kutokuwa na ujinga
katika aina hii]
☐ Cattle ☐ Goats ☐ Sheep
Other [Nyinginezo] [ ]

Do you think you would be more likely to sell animals that you know are infertile than those that are not [Je, unafikiri kuna uwezekano mkubwa kuuza wanyama ambayo ni tasa kuliko wale ambao sio tasa]?
Cattle [Ng'ombe] [ ]
Goats [Mbuzi] [ ]
Sheep [Kondoo] [ ]

9.39 Do you think you would be more likely to slaughter animals that you know are infertile than those that are not [Je, unafikiri kuna uwezekano mkubwa kuchinja wanyama ambayo ni tasa kuliko wale ambao sio tasa]?
Cattle [Ng'ombe] [ ]
Goats [Mbuzi] [ ]
Sheep [Kondoo] [ ]

9.40 What treatment do you give when an animal in your herd is infertile [Ni tiba gani unampa mnyama katika kundi lako ambayo ni tasa]? [ ]

9.41 Who do you normally go to for advice/treament when an animal in your herd is infertile [Kwa kawaida unakwenda kwa nani kupata ushauri/matibabu wakati mnyama wako ni tasa]?
☐ Neighbour [Jirani]
☐ Paravet [Paravet]
☐ Veterinarian [Bwana mifugo]
☐ Animal health assistant [Bwana afya msaidizi wa mifugo]
☐ Agrovet [Bwana kilimo na mifugo]
☐ Traditional healer [Mganga wa jadi]
☐ Spiritual/faith healer [Imani/mponyaji wa imani]
☐ Local livestock expert [Mtaalamu wa mifugo wa hapahapa]
☐ Other [Nyinginezo] [ ]

**SECTION 10: ANIMALS AROUND THE COMPOUND:**
We would now like to ask you a few questions about other animals that you may have seen around your compound, your village, or while grazing [Sasa hivi tunaweza kuuliza maswali kuhusu wanyama wengine walioko karibu na boma lako au ndani ya kijiji chako].

10.1 Have you seen rodents or evidence of rodents (e.g. faeces, urine, noises, rodent tracks, rodent damage) in or around your house in the past month [Je, umewahi kuwaona panya katika nyumba yako siku thelathini zilizopita]?
☐ Yes ☐ No

10.2 Do any members of this household do anything to control rodents [Je, yeyote wa wanakaya anafanya chochote kuzuia hawa panya]?
☐ Yes ☐ No

10.3 What type of rodent control do you use [Njia gani huwa unatumia]?

- ☐ Mechanical [Kuwatega, kuwapiga (mfano, mitego)]
- ☐ Chemical [Kutumia dawa/kemikali (mfano, sumu)]
- ☐ Biological [Kutumia njia za kibiologia (mfano, kufuga paka)]
- ☐ Other [Nyinginezo]

10.4 Do you see any of the following animals around your village during normal dry and wet seasons [Je, unaona wanyana wafuatao katika kijiji chako wakati wa kiangazi wa kawaida]?

| | | |
|---|---|---|
| Small monkeys [Karunguyeye] | ☐ Dry | ☐ Wet |
| Baboons [Kima] | ☐ Dry | ☐ Wet |
| Elephants [Tembo] | ☐ Dry | ☐ Wet |
| Zebra [Pundamilia] | ☐ Dry | ☐ Wet |
| Buffalo [Nyati / mbogo] | ☐ Dry | ☐ Wet |
| Antelope [Swala] | ☐ Dry | ☐ Wet |
| Giraffe [Twiga] | ☐ Dry | ☐ Wet |
| Wildebeest [Nyumbu] | ☐ Dry | ☐ Wet |
| Big cats (e.g. Lion, leopard) | ☐ Dry | ☐ Wet |
| Wild pigs (e.g bushpig, warthog) | ☐ Dry | ☐ Wet |
