## Supplementary figures and images for "The value of livestock abortion surveillance in Tanzania: identifying disease priorities and informing interventions"

### Supplementary Material 2

**Supplementary Material 2:**

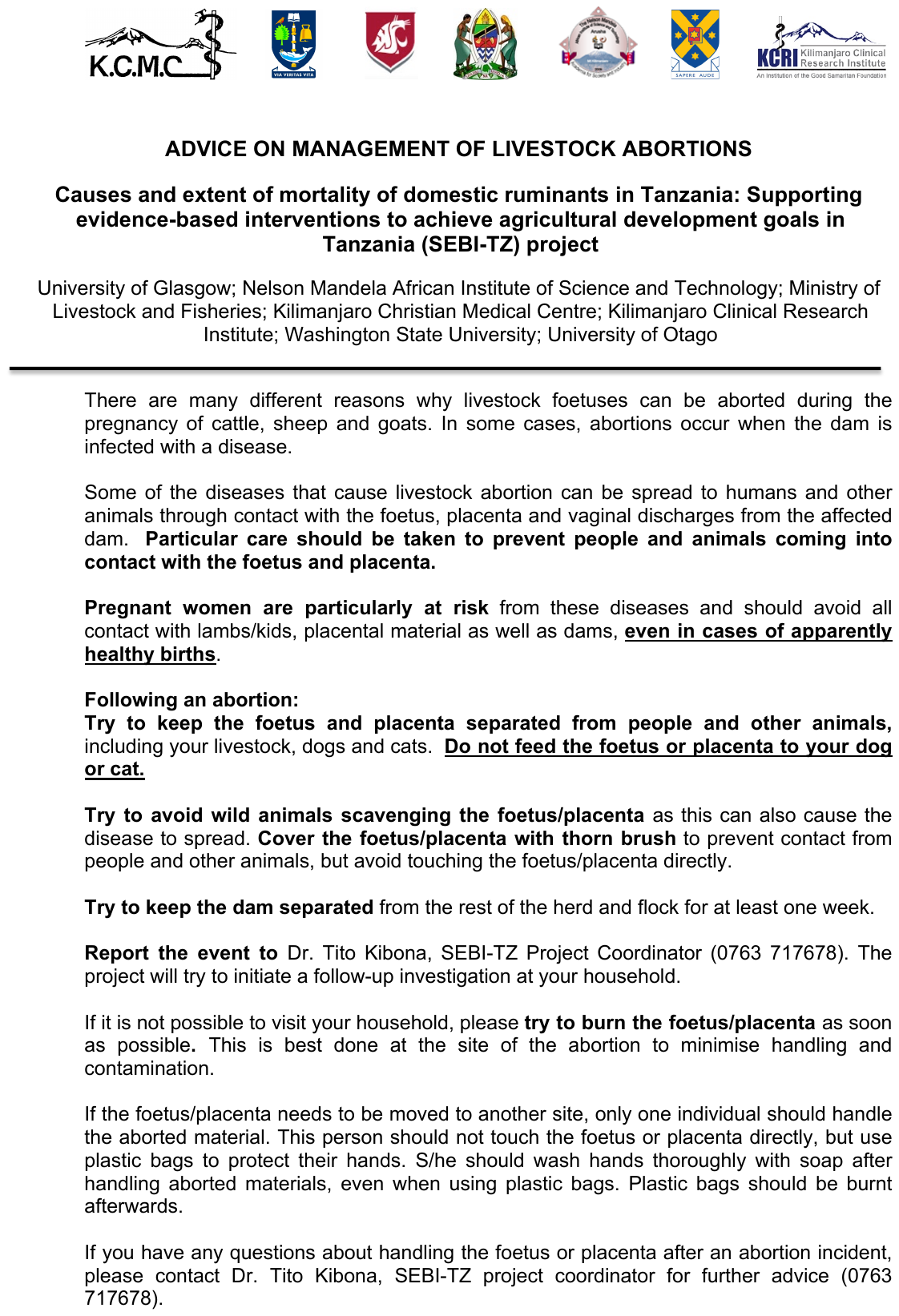
