## Supplementary Material 4 for "The value of livestock abortion surveillance in Tanzania: identifying disease priorities and informing interventions"

**Herd data**

The mean (median, range) number of cattle, goats and sheep per herd were 29.8 (7, 0 – 276), 55.9 (24, 0 – 817) and 47.5 (20, 0 – 1000), respectively, giving an approximate median herd composition ratio of 1 cattle to 3 goats to 3 sheep. The mean (median and range) number of adult female cattle, goats and sheep per herd were 17.5 9 (4 and 0 – 164), 35.4 (15 and 0 – 409) and 32.0 (12, and 0 – 800), respectively. The mean percentage of adult female cattle, goats and sheep per herd was 63.1%, 62.6% and 66.1%, respectively.
