## Supplementary Material 5 for "The value of livestock abortion surveillance in Tanzania: identifying disease priorities and informing interventions"

**Expected number of abortions per breed**

| **SPECIES** | **BREED** | **ACTUAL** | **EXPECTED** |
| --- | --- | --- | --- |
| Cattle | local | 16 | 61 |
| Goat | local | 77 | 94 |
| Sheep | local | 41 | 42 |
| Cattle | cross | 36 | 7 |
| Goat | cross | 17 | 2 |
| Sheep | cross | 3 | 2 |
| Cattle | exotic | 17 | 1 |
| Goat | exotic | 3 | 1 |
| Sheep | exotic | 0 | 0 |

The actual number of abortions reported for each species and breed and, based on the proportion of each breed in all the herds that reported cases, the expected number of abortions was calculated.
