## Supplementary Material 3a for "The value of livestock abortion surveillance in Tanzania: identifying disease priorities and informing interventions"

| TARAFA | WARD | VILLAGE | | | | N |
|---|---|---|---|---|---|---|
| | | | CATTLE | SHEEP | GOATS | PIGS |
| **Vunjo Mashariki** | Mwika kusini | Kimangaro | 1041 | 1334 | 350 | 1980 |
| | | Mawanjeni | 1260 | 1578 | 443 | 2210 |
| | | Kondeni | 988 | 1062 | 311 | 2300 |
| | | Matala | 1111 | 990 | 360 | 2400 |
| | | Kiruweni | 109 | 1053 | 123 | 1008 |
| **JUMLA** | | **5** | **5209** | **6017** | **1587** | **9898** |
| | Kirua vunjo kusini | Uchira | 1094 | 1600 | 307 | 102 |
| | | Mabungo | 1262 | 1360 | 310 | 90 |
| | | Yam Makaa | 998 | 1580 | 297 | 206 |
| | | Yamu | 866 | 970 | 320 | 232 |
| | | Uparo | 1249 | 1480 | 278 | 402 |
| **JUMLA** | | **5** | **5469** | **6990** | **1512** | **1032** |
| | Kilema kusini | Kilototoni | 769 | 1015 | 271 | 280 |
| | | Kilema Pofo | 849 | 957 | 190 | 292 |
| | | Masaera | 910 | 1183 | 182 | 181 |
| | | Leghomulo | 898 | 1177 | 213 | 177 |
| | | Kilema Chini | 800 | 882 | 200 | 283 |
| | | Marawe Kyura | 1058 | 897 | 271 | 465 |
| **JUMLA** | | **6** | **5282** | **6111** | **1266** | **1678** |
| | Kirua vunjo mashariki | Kileuo | 1191 | 1254 | 292 | 330 |
| | | Nganjoni | 1593 | 1262 | 470 | 321 |
| | | Mero | 933 | 1490 | 362 | 344 |
| | | Mrumeni | 1191 | 974 | 292 | 314 |
| **JUMLA** | | **4** | **4957** | **4980** | **1498** | **1309** |
| | Makuyuni | Makuyuni | 2464 | 2760 | 575 | 300 |
| | | Lotima | 2360 | 2614 | 600 | 295 |
| | | Himo | 1016 | 2608 | 510 | 292 |
| **JUMLA** | | **3** | 5840 | 7982 | 1685 | 887 |
| **Hai mashariki** | Arusha chini | Chemchem | 3925 | 4005 | 710 | 450 |
| | | Mikocheni | 4200 | 3865 | 890 | 440 |
| **JUMLA** | | **2** | **8125** | **7870** | **1600** | **890** |
| | Mabogini | Mtakuja | 1649 | 1794 | 500 | 207 |
| | | Mvuleni | 1559 | 1287 | 512 | 180 |
| | | Chekereni | 1876 | 1334 | 470 | 187 |
| | | Mabogini | 1511 | 1605 | 435 | 200 |
| **JUMLA** | | **4** | **6595** | **6020** | **1917** | **774** |
| | Kahe mashariki | Kiterini | 1067 | 1670 | 381 | 160 |
| | | Kyomu | 957 | 1432 | 420 | 188 |

| | | | | | | |
|---|---|---|---|---|---|---|
| | | Soko | 1365 | 1694 | 399 | 203 |
| | | Kochakindo | 1213 | 1810 | 277 | 211 |
| | | Ghona | 1152 | 2089 | 379 | 198 |
| **JUMLA** | | **5** | **5754** | **8695** | **1856** | **960** |
| | Kahe | Mwangaria | 1085 | 1581 | 492 | 140 |
| | | Ngasinyi | 1458 | 1600 | 300 | 170 |
| | | Kisangesageni | 1992 | 1500 | 285 | 166 |
| | | Mawala | 1849 | 1650 | 296 | 145 |
| | | Oria | 938 | 1236 | 301 | 159 |
| | | Rau river | 1387 | 1323 | 336 | 100 |
| **JUMLA** | | **6** | **8709** | **8890** | **2010** | **880** |
| **Kibosho** | Kindi | Kindi | 1508 | 1596 | 397 | 464 |
| | | Msasani | 1331 | 1416 | 311 | 312 |
| | | Sambarai | 1603 | 1317 | 365 | 298 |
| | | Chekereni weruweru | 1192 | 1658 | 370 | 360 |
| **JUMLA** | | **4** | **5634** | **5987** | **1443** | **1434** |
| **Kibosho** | Kirima | Boro | 1506 | 1379 | 490 | 560 |
| | | Kirima juu | 2117 | 1413 | 502 | 590 |
| | | Kirima kati | 1866 | 1047 | 479 | 517 |
| **JUMLA** | | **3** | **5489** | **3839** | **1471** | **1667** |
| | Okaoni | Omarini | 670 | 483 | 238 | 263 |
| | | Kitandu | 892 | 568 | 160 | 280 |
| | | Mloe | 582 | 417 | 140 | 286 |
| | | Sisamaro | 498 | 520 | 130 | 290 |
| | | Mkomilo | 502 | 442 | 142 | 203 |
| | | Dakau | 817 | 470 | 150 | 253 |
| **JUMLA** | | **6** | **3961** | **2900** | **960** | **1572** |
| | Kibosho magharibi | Manushi ndoo | 779 | 744 | 150 | 192 |
| | | Manushisinde | 712 | 654 | 170 | 222 |
| | | Kombo | 864 | 516 | 160 | 231 |
| | | Mkomongo | 661 | 602 | 152 | 182 |
| | | Kifuni | 679 | 760 | 140 | 189 |
| | | Umbwe sinde | 907 | 670 | 144 | 230 |
| | | Umbwe onana | 809 | 572 | 116 | 136 |
| **JUMLA** | | **7** | **5411** | **4518** | **1032** | **1382** |
| | Kibosho Mashariki | Mweka | 1195 | 1500 | 231 | 565 |
| | | Singa | 1900 | 1610 | 340 | 580 |
| | | Sungu | 1697 | 1230 | 328 | 543 |
| **JUMLA** | | **3** | **4792** | **4340** | **899** | **1688** |
| | Kibosho kati | Mauwa | 951 | 950 | 170 | 297 |

| | | | | | | |
|---|---|---|---|---|---|---|
| | | Uchau Kaskazini | 883 | 930 | 182 | 281 |
| | | Uchau Kusini | 1000 | 876 | 168 | 290 |
| | | Uri | 990 | 810 | 142 | 182 |
| | | Otaruni | 990 | 1134 | 188 | 386 |
| **JUMLA** | | **5** | **4752** | **4700** | **850** | **1436** |
| **Hai East** | Kimochi | Mowo | 1525 | 1005 | 455 | 325 |
| | | Shia | 1660 | 1010 | 312 | 193 |
| | | Mdawi | 1718 | 1211 | 410 | 275 |
| | | Sango | 1162 | 783 | 245 | 292 |
| **JUMLA** | | **4** | **6065** | **4009** | **1422** | **1085** |
| | Old Moshi magharibi | Tella | 1310 | 1533 | 480 | 480 |
| | | Mande | 1123 | 1628 | 492 | 430 |
| | | Mandaka mnono | 898 | 1424 | 443 | 460 |
| | | **3** | **3331** | **4585** | **1415** | **1370** |
| | Mbokomu | Korini juu | 1880 | 1470 | 460 | 670 |
| | | Korini kusini | 2007 | 1560 | 471 | 362 |
| | | Tema | 1940 | 1381 | 448 | 365 |
| **JUMLA** | | **3** | **5827** | **4411** | **1379** | **1397** |
| | Uru mashariki | Kishumundu | 578 | 565 | 190 | 202 |
| | | Mruiya | 497 | 610 | 207 | 179 |
| | | Mnini | 488 | 517 | 188 | 165 |
| | | Materuni | 612 | 477 | 210 | 152 |
| | | Mwasi kaskazini | 600 | 607 | 175 | 160 |
| | | Mwasi kusini. | 449 | 512 | 170 | 182 |
| | | Kyaseni | 578 | 599 | 220 | 170 |
| **JUMLA** | | **7** | **3761** | **3887** | **1360** | **1210** |
| | Uru Shimbwe | Shimbwe juu | 1497 | 1398 | 635 | 1000 |
| | | Shimbwe chini | 1693 | 1199 | 600 | 992 |
| **JUMLA** | | **2** | **3190** | **2597** | **1235** | **1992** |
| | Uru kusini | Okaseni | 464 | 677 | 180 | 286 |
| | | Kitandu | 563 | 620 | 167 | 170 |
| | | Longuo | 512 | 617 | 158 | 189 |
| | | Kimanganuni | 663 | 669 | 194 | 195 |
| | | Kariwa | 401 | 580 | 180 | 200 |
| | | Rau | 477 | 635 | 168 | 230 |
| **JUMLA** | | **6** | **3080** | **3798** | **1047** | **1270** |
| | Old moshi mashariki | Kidia | 778 | 1021 | 330 | 283 |

| | | | | | | |
|---|---|---|---|---|---|---|
| | | Tsuduni | 589 | 1317 | 323 | 293 |
| | | Kikarara | 708 | 983 | 190 | 271 |
| | | Mahoma | 695 | 764 | 321 | 296 |
| **JUMLA** | | **4** | **2770** | **4085** | **1164** | **1146** |
| | Uru kaskazini | Msuni | 890 | 745 | 196 | 320 |
| | | Ongoma | 792 | 790 | 215 | 298 |
| | | Mrawi | 904 | 680 | 200 | 301 |
| | | Njari | 936 | 754 | 176 | 341 |
| **JUMLA** | | **4** | **3522** | **2969** | **787** | **1260** |
| **Vunjo Mashariki** | Kilema Kaskazini | Makame juu | 1292 | 1307 | 237 | 472 |
| | | Makame chini | 1389 | 1410 | 189 | 277 |
| | | Kyou | 1190 | 1385 | 214 | 380 |
| | | Ruwa | 1250 | 889 | 220 | 392 |
| **JUMLA** | **JUMLA** | **4** | **5121** | **4991** | **860** | **1521** |
| | Kilema kati | Kimaroroni | 1030 | 1300 | 232 | 494 |
| | | Ngangu | 1110 | 1400 | 259 | 348 |
| | | Mkyashi | 1010 | 1310 | 211 | 383 |
| | | Rosho | 967 | 710 | 188 | 464 |
| **JUMLA** | | **4** | **4117** | **4800** | **890** | **1689** |
| | Mwika Kaskazini | Lole Marera | 956 | 907 | 406 | 346 |
| | | Maring'a | 910 | 1272 | 182 | 283 |
| | | Msae Nganyeni | 870 | 976 | 197 | 292 |
| | | Mrimbo uuwo | 797 | 821 | 255 | 311 |
| | | Msae Kinyamvuo | 1195 | 1012 | 260 | 320 |
| **JUMLA** | | **5** | **4728** | **4988** | **1300** | **1552** |
| | Marangu magharibi | Kitowo | 768 | 824 | 140 | 251 |
| | | Kiraracha | 790 | 704 | 137 | 260 |
| | | Komalyangoe | 902 | 710 | 122 | 190 |
| | | Kyala | 577 | 677 | 162 | 282 |
| | | Nduweni | 560 | 661 | 150 | 170 |
| | | Komela | 522 | 702 | 143 | 230 |
| | | Mbahe | 1192 | 639 | 116 | 307 |
| **JUMLA** | | **7** | **5312** | **4917** | **970** | **1690** |
| | Marangu Mashariki | Sembeti | 650 | 850 | 114 | 187 |
| | | Lyamrakana | 885 | 905 | 170 | 168 |
| | | Arisi | 990 | 712 | 167 | 150 |
| | | Lyasongoro | 918 | 767 | 155 | 149 |
| | | Samanga | 1007 | 790 | 201 | 158 |
| | | Mshiri | 792 | 869 | 213 | 170 |

| | | | | | | |
|---|---|---|---|---|---|---|
| | | Rauya | 756 | 1051 | 114 | 260 |
| **JUMLA** | | **7** | **5998** | **5944** | **1200** | **1242** |
| | Mamba kaskazini | Mboni | 1289 | 1426 | 235 | 248 |
| | | Komakundi | 1391 | 1502 | 212 | 260 |
| | | Kokirie | 991 | 1517 | 196 | 271 |
| | | Kotela | 1086 | 1221 | 227 | 193 |
| **JUMLA** | | **4** | **4757** | **5666** | **870** | **972** |
| | Mamba kusini | Lekura | 685 | 671 | 140 | 320 |
| | | Kiria | 702 | 516 | 150 | 309 |
| | | Mkolowoni | 710 | 580 | 155 | 292 |
| | | Kimangara | 590 | 605 | 161 | 384 |
| | | Kimbogho | 791 | 687 | 108 | 316 |
| **JUMLA** | | **5** | **3478** | **3059** | **714** | **1521** |
| **Vunjo magharibi** | Kirua vunjo Magharibi | Nduoni | 687 | 914 | 180 | 230 |
| | | Maruwa | 690 | 811 | 192 | 249 |
| | | Kanango | 698 | 620 | 202 | 270 |
| | | Kanji | 600 | 582 | 213 | 185 |
| | | Kwamare | 712 | 577 | 179 | 190 |
| | | Manu | 563 | 694 | 181 | 197 |
| | | Iwa | 796 | 783 | 143 | 270 |
| **JUMLA** | | **7** | **4746** | **4981** | **1290** | **1591** |
| **JUMLA KUU** | | **145** | **162782** | **159,326** | **39,691** | **40,586** |

| NUMBER OF LIVESTOCK | | | | |
|---|---|---|---|---|
| RABBITS | CHICKENS | DOGS | CATS | DONKEYS |
| | 6380 | | | |
| - | 5760 | | | |
| - | 5997 | | | |
| - | 6013 | | | |
| - | 7734 | | | |
| **--** | **31884** | **890** | 392 | - |
| - | 5866 | | | |
| - | 5004 | | | |
| - | 5997 | | | |
| - | 5700 | | | |
| - | 6713 | | | |
| **-** | **29280** | **1120** | **600** | **-** |
| - | 5700 | | | |
| 226 | 6910 | | | |
| 320 | 6279 | | | |
| - | 4912 | | | |
| 262 | 3678 | | | |
| - | 6621 | | | |
| **908** | **34100** | **1300** | 654 | - |
| - | 8439 | | | |
| - | 6980 | | | |
| - | 7665 | | | |
| - | 7828 | | | |
| **-** | **30912** | **800** | 551 | - |
| - | 9220 | | | |
| - | 10120 | | | |
| - | 9360 | | | |
| - | 28700 | 1320 | 364 | 93 |
| - | 13900 | | | |
| - | 12302 | | | |
| **-** | **26202** | 1676 | 624 | 239 |
| - | 6712 | | | |
| | 6010 | | | |
| | 5849 | | | |
| - | 6138 | | | |
| **-** | **24709** | **1400** | 265 | 210 |
| - | 6220 | | | |
| - | 5159 | | | |

| | | | | |
|---|---|---|---|---|
| - | 5285 | | | |
| - | 6187 | | | |
| - | 7249 | | | |
| **-** | **30100** | **1054** | 321 | 197 |
| - | 5800 | | | |
| - | 5669 | | | |
| - | 4914 | | | |
| - | 4800 | | | |
| - | 5400 | | | |
| - | 4617 | | | |
| **-** | **31200** | **1765** | 426 | 162 |
| - | 8560 | | | |
| 200 | 8109 | | | |
| 300 | 7917 | | | |
| 118 | 7624 | | | |
| **618** | **32210** | **790** | 332 | - |
| - | 12400 | | | |
| - | 12194 | | | |
| - | 9106 | | | |
| **-** | **33700** | **650** | 219 | - |
| - | 4910 | | | |
| - | 4872 | | | |
| - | 4511 | | | |
| - | 7966 | | | |
| - | 5793 | | | |
| - | 5124 | | | |
| **-** | **33176** | **640** | 376 | - |
| - | 4800 | | | |
| - | 4212 | | | |
| - | 3870 | | | |
| - | 3700 | | | |
| - | 3655 | | | |
| - | 4014 | | | |
| - | 5948 | | | |
| **-** | **30199** | **890** | 470 | - |
| - | 10344 | | | |
| - | 11990 | | | |
| - | 12666 | | | |
| **-** | **35000** | **1230** | 548 | - |
| | 8219 | | | |

| | | | | |
|---|---|---|---|---|
| - | 7768 | | | |
| - | 6912 | | | |
| - | 7690 | | | |
| - | 8163 | | | |
| **-** | **38752** | **1322** | 380 | - |
| 186 | 9398 | | | |
| 234 | 9912 | | | |
| 180 | 8170 | | | |
| 89 | 9398 | | | |
| **689** | **37000** | **977** | 453 | - |
| 700 | 13560 | | | |
| 589 | 14100 | | | |
| - | 12259 | | | |
| **1289** | **39917** | **644** | 380 | - |
| 320 | 14033 | | | |
| 289 | 11818 | | | |
| 218 | 12926 | | | |
| **827** | **38777** | **768** | 429 | - |
| 254 | 3742 | | | |
| 360 | 3847 | | | |
| 466 | 2852 | | | |
| - | 3170 | | | |
| - | 3510 | | | |
| - | 3776 | | | |
| - | 6223 | | 313 | - |
| **1080** | **27120** | **428** | | |
| - | 16800 | | | |
| - | 15500 | | | |
| **-** | **32300** | **598** | 388 | - |
| 540 | 5648 | | | |
| 670 | 6200 | | | |
| 680 | 4313 | | | |
| 310 | 4800 | | | |
| - | 5515 | | | |
| - | 6324 | | | |
| **2200** | **32800** | **888** | 576 | - |
| | 6275 | | | |

| | | | | |
|---|---|---|---|---|
| 359 | 6527 | | | |
| 200 | 5810 | | | |
| 470 | 7497 | | | |
| **1029** | **26109** | **676** | 445 | - |
| - | 7667 | | | |
| - | 7000 | | | |
| - | 6916 | | | |
| - | 5084 | | | |
| **-** | **26667** | **810** | 300 | - |
| 900 | 6335 | | | |
| 800 | 8901 | | | |
| 650 | 8780 | | | |
| 668 | 9307 | | | |
| **3018** | **33323** | **390** | 661 | - |
| - | 8594 | | | |
| - | 7910 | | | |
| - | 7760 | | | |
| - | 6112 | | | |
| **-** | **30376** | **298** | **432** | - |
| 400 | 7218 | | | |
| 600 | 4660 | | | |
| - | 6718 | | | |
| - | 6800 | | | |
| - | 5613 | | | |
| **1000** | **31009** | **810** | **466** | - |
| - | 6579 | | | |
| - | 6720 | | | |
| - | 5440 | | | |
| - | 5373 | | | |
| - | 5110 | | | |
| - | 4990 | | | |
| - | 4811 | | | |
| **-** | **39021** | **706** | 410 | - |
| 500 | 4410 | | | |
| - | 4222 | | | |
| 412 | 4943 | | | |
| - | 4823 | | | |
| - | 4337 | | | |
| - | 4500 | | | |

| | | | | |
|---|---|---|---|---|
| - | 3635 | | | |
| **912** | **30870** | **752** | 309 | - |
| - | 6845 | | | |
| - | 6260 | | | |
| - | 6172 | | | |
| - | 7703 | | | |
| **-** | **26980** | **360** | 259 | - |
| - | 6450 | | | |
| 560 | 6132 | | | |
| 330 | 5790 | | | |
| - | 5610 | | | |
| - | 7245 | | | |
| **890** | **31227** | **289** | 388 | - |
| - | 5606 | | | |
| - | 4319 | | | |
| - | 4170 | | | |
| - | 4267 | | | |
| - | 3978 | | | |
| - | 3810 | | | |
| - | 5534 | 471 | 350 | - |
| **-** | **31684** | | | |
| **14,460** | **985,304** | **26,712** | **13,390** | **901** |

| Region | District | Ward | Village | Cow | Goat | Sheep |
|---|---|---|---|---|---|---|
| Arusha | Monduli | Lepurko | Lepurko Shul | 3278 | 3633 | 2154 |
| Arusha | Monduli | Lepurko | Eng'arooji | 2822 | 3215 | 1675 |
| Arusha | Monduli | Lepurko | Mti mmoja | 4766 | 3548 | 2179 |
| Arusha | Monduli | Lepurko | Losimingori | 7938 | 2702 | 3266 |
| Arusha | Monduli | Lepurko | Nanja | 2564 | 3688 | 1688 |
| Arusha | Monduli | Lepurko | | 21368 | 16786 | 10962 |
| Arusha | Monduli | Meserani | Meserani juu | 4751 | 3966 | 3452 |
| Arusha | Monduli | Meserani | Naaarami | 4643 | 3783 | 2311 |
| Arusha | Monduli | Meserani | Bwawani | 1300 | 2738 | 1265 |
| Arusha | Monduli | Meserani | | 10694 | 10487 | 7028 |
| Arusha | Monduli | Sepeko | Arkaria | 3628 | 2189 | 2189 |
| Arusha | Monduli | Sepeko | Arkatan | 1461 | 2534 | 1534 |
| Arusha | Monduli | Sepeko | Lashaine | 1814 | 2328 | 1328 |
| Arusha | Monduli | Sepeko | Lendikinya | 1089 | 3377 | 1377 |
| Arusha | Monduli | Sepeko | | 7992 | 10428 | 6428 |
| Arusha | Monduli | Monduli Juu | Eluway | 1976 | 3276 | 1276 |
| Arusha | Monduli | Monduli Juu | Enguik | 2014 | 2643 | 1643 |
| Arusha | Monduli | Monduli Juu | Mfereji | 2376 | 3274 | 1274 |
| Arusha | Monduli | Monduli Juu | Idonyonado | 1264 | 3193 | 1193 |
| Arusha | Monduli | Monduli Juu | Emairete | 3412 | 4215 | 1215 |
| Arusha | Monduli | Monduli Juu | | 11042 | 16601 | 6601 |
| Arusha | Monduli | Moita | Kilimatinde | 2666 | 2286 | 1286 |
| Arusha | Monduli | Moita | Moita Kilorit | 2388 | 2168 | 1168 |
| Arusha | Monduli | Moita | Moita Bwawa | 2392 | 2173 | 1173 |
| Arusha | Monduli | Moita | Moita Kipok | 2744 | 1766 | 1766 |
| Arusha | Monduli | Moita | | 10190 | 8393 | 5393 |
| Arusha | Monduli | Engutoto | Olarash | 1538 | 844 | 893 |
| Arusha | Monduli | Engutoto | Ngarash | 1645 | 1055 | 1055 |
| Arusha | Monduli | Engutoto | Mlimani | 1566 | 654 | 654 |
| Arusha | Monduli | Engutoto | | 4749 | 2553 | 2602 |
| Arusha | Monduli | Monduli Mjin | TMA & SOF | 0 | 459 | 276 |
| Arusha | Monduli | Monduli Mjin | Monduli Mjin | 168 | 132 | 83 |
| Arusha | Monduli | Monduli Mjini | | 168 | 591 | 359 |
| Arusha | Monduli | Lolkisale | Tukusi | 2371 | 2645 | 2645 |
| Arusha | Monduli | Lolkisale | Lolkisale | 7342 | 5489 | 5489 |
| Arusha | Monduli | Lolkisale | NAFCO | 3415 | 2177 | 2177 |
| Arusha | Monduli | Lolkisale | Lemoti | 10432 | 7484 | 7484 |
| Arusha | Monduli | Lolkisale | | 23560 | 17795 | 17795 |
| Arusha | Monduli | Makuyuni | Naiti | 10122 | 6593 | 6593 |
| Arusha | Monduli | Makuyuni | Makuyuni | 9878 | 7985 | 7285 |
| Arusha | Monduli | Makuyuni | Mbuyuni | 4399 | 4262 | 4262 |
| Arusha | Monduli | Makuyuni | | 24399 | 18840 | 18140 |
| Arusha | Monduli | Mswakini | Mswakini Juu | 8675 | 5487 | 5487 |
| Arusha | Monduli | Mswakini | Naitolia | 3854 | 3126 | 3126 |
| Arusha | Monduli | Mswakini | Mswakini Ch | 6965 | 4385 | 4385 |

| | | | | | | |
|---|---|---|---|---|---|---|
| Arusha | Monduli | Mswakini | | 19494 | 12998 | 12998 |
| Arusha | Monduli | Esilalei | Oltukai | 10766 | 8471 | 8699 |
| Arusha | Monduli | Esilalei | Mungere | 2744 | 3512 | 1512 |
| Arusha | Monduli | Esilalei | Losirwa | 10548 | 7132 | 7132 |
| Arusha | Monduli | Esilalei | Manyara ranc | 1472 | 0 | 677 |
| Arusha | Monduli | Esilalei | Esilalei | 11538 | 7633 | 7186 |
| Arusha | Monduli | Esilalei | | 37068 | 26748 | 25206 |
| Arusha | Monduli | Selela | Mbaash | 11584 | 10871 | 12532 |
| Arusha | Monduli | Selela | Selela | 11499 | 10266 | 13422 |
| Arusha | Monduli | Selela | | 23083 | 21137 | 25954 |
| Arusha | Monduli | Engaruka | Engaruka Juu | 8932 | 8326 | 9833 |
| Arusha | Monduli | Engaruka | Engaruka Chi | 7744 | 7286 | 8468 |
| Arusha | Monduli | Engaruka | | 16676 | 15612 | 18301 |
| Arusha | Monduli | Mto wa Mbu | Barabarani | 1032 | 572 | 642 |
| Arusha | Monduli | Mto wa Mbu | Migungani | 2422 | 948 | 855 |
| Arusha | Monduli | Mto wa Mbu | Jangwani | 1488 | 1329 | 1012 |
| Arusha | Monduli | Mto wa Mbu | | 4942 | 2849 | 2509 |
| Arusha | Monduli | Majengo | Majengo | 488 | 1732 | 733 |
| Arusha | Monduli | Majengo | Migombani | 386 | 583 | 214 |
| Arusha | Monduli | Majengo | Kigongoni | 638 | 1528 | 547 |
| Arusha | Monduli | Majengo | Migungani | 473 | 1064 | 694 |
| Arusha | Monduli | Majengo | | 1985 | 4907 | 2188 |
| Arusha | Monduli | | JUMLA WII | 217410 | 186725 | 162464 |

| Dog | Pig |
|---|---|
| 126 | 0 |
| 87 | 0 |
| 129 | 0 |
| 186 | 0 |
| 326 | 0 |
| 854 | 0 |
| 388 | 0 |
| 124 | 0 |
| 144 | 0 |
| 656 | 0 |
| 125 | 0 |
| 93 | 16 |
| 137 | 0 |
| 114 | 0 |
| 469 | 16 |
| 149 | 0 |
| 176 | 13 |
| 127 | 0 |
| 113 | 0 |
| 278 | 26 |
| 843 | 39 |
| 118 | 0 |
| 192 | 0 |
| 134 | 24 |
| 112 | 0 |
| 556 | 24 |
| 208 | 54 |
| 195 | 38 |
| 244 | 49 |
| 647 | 141 |
| 198 | 71 |
| 188 | 122 |
| 386 | 193 |
| 99 | 0 |
| 115 | 47 |
| 123 | 472 |
| 108 | 0 |
| 445 | 519 |
| 149 | 0 |
| 323 | 79 |
| 274 | 0 |
| 746 | 79 |
| 263 | 0 |
| 264 | 0 |
| 236 | 0 |

| | |
|---|---|
| 763 | 0 |
| 123 | 0 |
| 0 | 0 |
| 236 | 22 |
| 17 | 0 |
| 163 | 0 |
| 539 | 22 |
| 168 | 0 |
| 213 | 0 |
| 381 | 0 |
| 157 | 0 |
| 188 | 0 |
| 345 | 0 |
| 186 | 69 |
| 0 | 0 |
| 194 | 38 |
| 380 | 107 |
| 204 | 123 |
| 164 | 144 |
| 141 | 54 |
| 197 | 0 |
| 706 | 321 |
| 8716 | 1461 |

| Region | District | Ward | Village | Cow | Goat | Sheep |
|---|---|---|---|---|---|---|
| Arusha | Karatu | Baray | Qandend | 5514 | 13392 | 1536 |
| Arusha | Karatu | Baray | Mbuga Nyeku | 7532 | 7983 | 1848 |
| Arusha | Karatu | Baray | Dumbechand | 13849 | 4716 | 2013 |
| Arusha | Karatu | Baray | Jobaj | 4279 | 8696 | 1273 |
| Arusha | Karatu | Baray | Matala | 18194 | 10892 | 1655 |
| Arusha | Karatu | Qurus | Bashay | 6714 | 5953 | 1302 |
| Arusha | Karatu | Qurus | Gongali | 4466 | 4662 | 1113 |
| Arusha | Karatu | Qurus | Qurus | 6243 | 5560 | 882 |
| Arusha | Karatu | Daa | Changarawe | 2737 | 2970 | 232 |
| Arusha | Karatu | Daa | Endashagwet | 4483 | 10052 | 1871 |
| Arusha | Karatu | Daa | Mang'ola Juu | 8381 | 6594 | 1773 |
| Arusha | Karatu | Daa | Makhoromba | 7750 | 3888 | 1734 |
| Arusha | Karatu | Mang'ola | Mang'ola Ba | 5550 | 2764 | 2347 |
| Arusha | Karatu | Mang'ola | Maleckchand | 5299 | 2114 | 2261 |
| Arusha | Karatu | Mang'ola | Endamaghan | 7812 | 11284 | 6055 |
| Arusha | Karatu | Mang'ola | Laghangarer | 5349 | 5250 | 4109 |
| Arusha | Karatu | Endamarariek | Bassodawish | 7601 | 6346 | 2887 |
| Arusha | Karatu | Endamarariek | Endamarariek | 5021 | 8542 | 2109 |
| Arusha | Karatu | Endamarariek | Endallah | 2580 | 9712 | 725 |
| Arusha | Karatu | Endamarariek | Getamock | 10290 | 8908 | 2395 |
| Arusha | Karatu | Endamarariek | Khusumay | 4059 | 5118 | 1625 |
| Arusha | Karatu | Kansay | Kansay | 8911 | 3754 | 1459 |
| Arusha | Karatu | Kansay | Ngaibara | 2245 | 1920 | 533 |
| Arusha | Karatu | Kansay | Laja | 11208 | 4624 | 1913 |
| Arusha | Karatu | Kansay | Kambi Faru | 6254 | 4252 | 1053 |
| Arusha | Karatu | Endabash | Endabash | 6353 | 8080 | 2341 |
| Arusha | Karatu | Endabash | Qaru | 8479 | 9022 | 4375 |
| Arusha | Karatu | Buger | Buger | 11234 | 6722 | 1423 |
| Arusha | Karatu | Buger | Endanyowet | 6935 | 5012 | 1043 |
| Arusha | Karatu | Buger | Ayalio | 4895 | 4462 | 1293 |
| Arusha | Karatu | Mbulumbulu | Lositete | 5320 | 5552 | 1893 |
| Arusha | Karatu | Mbulumbulu | Upper Kitete | 3772 | 5086 | 1353 |
| Arusha | Karatu | Mbulumbulu | Slahhamo | 3676 | 4692 | 1313 |
| Arusha | Karatu | Mbulumbulu | Kambi Simba | 4229 | 7320 | 1309 |
| Arusha | Karatu | Rhotia | Rhotia Kati | 3835 | 3300 | 1713 |
| Arusha | Karatu | Rhotia | Rhotia Kaina | 4004 | 3952 | 1275 |
| Arusha | Karatu | Rhotia | Kilimatembo | 3184 | 3526 | 1023 |
| Arusha | Karatu | Rhotia | Kilimamoja | 4406 | 5012 | 2053 |
| Arusha | Karatu | Rhotia | Chemchem | 4405 | 5134 | 1593 |
| Arusha | Karatu | Ganako | Ayalabe | 3602 | 2104 | 979 |
| Arusha | Karatu | Ganako | Tloma | 4075 | 3492 | 1345 |
| Arusha | Karatu | Oldeani | Oldeani | 3427 | 5160 | 1795 |
| Arusha | Karatu | Karatu | G/arusha | 4245 | 2312 | 1203 |
| Arusha | Karatu | Karatu | Karatu mjini | 245 | 98 | 32 |
| Arusha | Karatu | Karatu | G/lambo | 5201 | 2012 | 2412 |

| | | | | | | |
|---|---|---|---|---|---|---|
| Arusha | Karatu | Jumla kuu | | 267,843 | 251,996 | 78,469 |

| Pig | Dog |
|---|---|
| 425 | 449 |
| 330 | 332 |
| 215 | 298 |
| 110 | 132 |
| 34 | 289 |
| 535 | 196 |
| 408 | 245 |
| 302 | 247 |
| 261 | 110 |
| 212 | 30 |
| 62 | 226 |
| 118 | 174 |
| 234 | 35 |
| 45 | 27 |
| 35 | 40 |
| 37 | 36 |
| 356 | 488 |
| 419 | 478 |
| 415 | 543 |
| 394 | 546 |
| 165 | 689 |
| 418 | 35 |
| 131 | 38 |
| 182 | 19 |
| 313 | 48 |
| 1,261 | 955 |
| 735 | 356 |
| 259 | 187 |
| 205 | 198 |
| 257 | 148 |
| 106 | 86 |
| 169 | 98 |
| 354 | 56 |
| 186 | 81 |
| 205 | 216 |
| 258 | 229 |
| 235 | 108 |
| 126 | 102 |
| 218 | 182 |
| 238 | 198 |
| 186 | 161 |
| 246 | 256 |
| 312 | 234 |
| 87 | 521 |
| 123 | 189 |

| | |
|---|---|
| 11,922 | 10,311 |

| Region | District | Ward | Imported Cattle | Local Cattle | Total Cattle |
|---|---|---|---|---|---|
| Manyara | Babati Rural | Gidas | | 6760 | 6760 |
| Manyara | Babati Rural | Mwada | | 25391 | 25391 |
| Manyara | Babati Rural | Ufana | 80 | 6880 | 6960 |
| Manyara | Babati Rural | Dabil | 91 | 15159 | 15250 |
| Manyara | Babati Rural | Magara | 18 | 10220 | 10238 |
| Manyara | Babati Rural | Duru | 0 | 7284 | 7284 |
| Manyara | Babati Rural | Endakiso | 61 | 4785 | 4846 |
| Manyara | Babati Rural | Madunga | 0 | 12174 | 12174 |
| Manyara | Babati Rural | Riroda | 421 | 4716 | 5137 |
| Manyara | Babati Rural | Dareda | 519 | 6984 | 7503 |
| Manyara | Babati Rural | Arri | 0 | 5625 | 5625 |
| Manyara | Babati Rural | Ayasanda | 73 | 2968 | 3041 |
| Manyara | Babati Rural | Magugu | 381 | 11670 | 12051 |
| Manyara | Babati Rural | Bashneti | 110 | 7661 | 7771 |
| Manyara | Babati Rural | Nkaiti | 35 | 38827 | 38862 |
| Manyara | Babati Rural | Boay | | | |
| Manyara | Babati Rural | Secheda | 65 | 7722 | 7787 |
| Manyara | Babati Rural | Kiru | 0 | 5813 | 5813 |
| Manyara | Babati Rural | Gallapo | 401 | 17384 | 17785 |
| Manyara | Babati Rural | Qash | 382 | 13641 | 14023 |
| Manyara | Babati Rural | Mamire | 68 | 8529 | 8597 |
| Manyara | Kiteto | Partimbo | | | 14954 |
| Manyara | Kiteto | Namelok | | | 17889 |
| Manyara | Kiteto | Bwagamoyo | | | 1233 |
| Manyara | Kiteto | Kibaya | | | 0 |
| Manyara | Kiteto | Kijungu | | | 6686 |
| Manyara | Kiteto | Lengatei | | | 32096 |
| Manyara | Kiteto | Loolera | | | 11103 |
| Manyara | Kiteto | Engusero | | | 9121 |
| Manyara | Kiteto | Matui | | | 13056 |
| Manyara | Kiteto | Dosidosi | | | 7472 |
| Manyara | Kiteto | Songambele | | | 20578 |
| Manyara | Kiteto | Magungu | | | 14169 |
| Manyara | Kiteto | Makami | | | 35018 |
| Manyara | Kiteto | Ndedo | | | 27561 |
| Manyara | Kiteto | Njoro | | | 19235 |
| Manyara | Kiteto | Kiperesa | | | 17521 |
| Manyara | Kiteto | Sunya | | | 37102 |
| Manyara | Kiteto | Dongo | | | 30337 |
| Manyara | Hanang | Bassotu | | | 38,409 |
| Manyara | Hanang | Mulbadaw | | | 10,294 |
| Manyara | Hanang | Hirbadaw | | | 16,350 |
| Manyara | Hanang | Getanuwas | | | 8,524 |
| Manyara | Hanang | Laghanga | | | 9,825 |
| Manyara | Hanang | Garawja | | | 6,501 |

| | | | | | |
|---|---|---|---|---|---|
| Manyara | Hanang | Bassodesh | | | 10,167 |
| Manyara | Hanang | Measkron | | | 6,750 |
| Manyara | Hanang | Endagaw | | | 2,304 |
| Manyara | Hanang | Endasiwold | | | 2,824 |
| Manyara | Hanang | Endasak | | | 892 |
| Manyara | Hanang | Gitting | | | 11,476 |
| Manyara | Hanang | Masakta | | | 4,314 |
| Manyara | Hanang | Masqaroda | | | 16,377 |
| Manyara | Hanang | G’babieg | | | 3,639 |
| Manyara | Hanang | B’lalu | | | 16,572 |
| Manyara | Hanang | Lalaji | | | 17,767 |
| Manyara | Hanang | Gehandu | | | 23,367 |
| Manyara | Hanang | Ishponga | | | 6,007 |
| Manyara | Hanang | Simbay | | | 5,870 |
| Manyara | Hanang | Sirop | | | 5,540 |
| Manyara | Hanang | Hidet | | | 3,362 |
| Manyara | Hanang | G’lang | | | 9,038 |
| Manyara | Hanang | Katesh | | | 874 |
| Manyara | Hanang | Ganana | | | 190 |
| Manyara | Hanang | Jorodom | | | 1,625 |
| Manyara | Hanang | Dumbeta | | | 3,560 |
| Manyara | Hanang | Nangwa | | | 2,539 |
| Manyara | Hanang | Dirma | | | 18,220 |
| Manyara | Hanang | Wareta | | | 6,232 |
| Manyara | Hanang | Dawari | | | 3,384 |
| Manyara | Hanang | Gendabi | | | 4,960 |
| Manyara | Hanang | Mogitu | | | 8,459 |
| Manyara | Babati Urban | Babati | 1448 | 1451 | 1451 |
| Manyara | Babati Urban | Bagara | 224 | 3750 | 3750 |
| Manyara | Babati Urban | Bonga | 560 | 2215 | 2215 |
| Manyara | Babati Urban | Maisaka | 55 | 3627 | 3627 |
| Manyara | Babati Urban | Mutuka | 5 | 3027 | 3027 |
| Manyara | Babati Urban | Nangara | 693 | 2093 | 2093 |
| Manyara | Babati Urban | Sigino | 193 | 4620 | 4620 |
| Manyara | Babati Urban | Singe | 217 | 1670 | 1670 |
| Manyara | Simanjiro | Naberara | | | 80563 |
| Manyara | Simanjiro | Komolo | | | 12043 |
| Manyara | Simanjiro | Ngorika | | | 8486 |
| Manyara | Simanjiro | R/remit | | | 24465 |
| Manyara | Simanjiro | Shambarai | | | 12684 |
| Manyara | Simanjiro | M/tembo | | | 9763 |
| Manyara | Simanjiro | Endiamtu | | | 515 |
| Manyara | Simanjiro | Mererani | | | 170 |
| Manyara | Simanjiro | Terrat | | | 25391 |
| Manyara | Simanjiro | Kitwai | | | 28892 |
| Manyara | Simanjiro | Langai | | | 17989 |

| | | | | | |
|---|---|---|---|---|---|
| Manyara | Simanjiro | Orksement | | | 2280 |
| Manyara | Simanjiro | Oljro No 5 | | | 20164 |
| Manyara | Simanjiro | Loboisorot | | | 38559 |
| Manyara | Simanjiro | Emboreet | | | 34263 |
| Manyara | Simanjiro | Naisinyai | | | 29915 |
| Manyara | Simanjiro | Loibosoit | | | 18145 |
| Manyara | Simanjiro | Endonyongijape | | | 16517 |

| Imported Goats | Local Goats | Total Goats | Sheep | Pigs | Dogs |
|---|---|---|---|---|---|
| | 4881 | 4881 | 2249 | 163 | 716 |
| | 16642 | 16642 | 7493 | 174 | 778 |
| 22 | 7469 | 7491 | 329 | 217 | 797 |
| 17 | 7731 | 7748 | 5354 | 611 | 1322 |
| 55 | 6656 | 6711 | 1689 | 618 | 837 |
| | 5150 | 5150 | 1834 | 131 | 889 |
| 47 | 7141 | 7188 | 3301 | 29 | 473 |
| 0 | 2328 | 2328 | 2766 | 460 | 488 |
| 73 | 5781 | 5854 | 1367 | 396 | 845 |
| 223 | 4442 | 4665 | 2398 | 1475 | 743 |
| 0 | 5236 | 5236 | 1933 | 464 | 526 |
| 1 | 3290 | 3291 | 1627 | 9 | 447 |
| 51 | 11847 | 11898 | 4839 | 620 | 1173 |
| 89 | 4500 | 4589 | 3583 | 2667 | 410 |
| 74 | 21947 | 22021 | 14871 | 22 | 2687 |
| 46 | 6843 | 6889 | 447 | 507 | 600 |
| 0 | 6005 | 6005 | 1724 | 459 | 727 |
| 163 | 16340 | 16503 | 4453 | 214 | 1521 |
| 100 | 9044 | 9144 | 3799 | 47 | 1136 |
| 79 | 11673 | 11752 | 2116 | 91 | 450 |
| | | 9494 | 2226 | 9 | 319 |
| | | 30434 | 5541 | 159 | 580 |
| | | 1112 | 16 | 174 | 111 |
| | | 489 | 0 | 47 | 32 |
| | | 4231 | 1476 | 0 | 78 |
| | | 13641 | 3757 | 0 | 666 |
| | | 19590 | 9605 | 0 | 164 |
| | | 5694 | 6181 | 342 | 478 |
| | | 5396 | 1024 | 329 | 451 |
| | | 2879 | 388 | 279 | 158 |
| | | 14774 | 3215 | 0 | 126 |
| | | 15613 | 6355 | 149 | 107 |
| | | 27960 | 15329 | 0 | 136 |
| | | 6850 | 3774 | 0 | 117 |
| | | 14524 | 5001 | 7 | 383 |
| | | 15474 | 5627 | 0 | 353 |
| | | 18614 | 11200 | 17 | 207 |
| | | 19152 | 1682 | 414 | 325 |
| | | 15,360 | 2,234 | 824 | 1,650 |
| | | 7,817 | 3,874 | 69 | 133 |
| | | 13,580 | 900 | 210 | 393 |
| | | 5,435 | 2,337 | 236 | 807 |
| | | 7,564 | 5764 | 364 | 1,482 |
| | | 7,327 | 3,111 | 780 | 612 |

| | | | | | |
|---|---|---|---|---|---|
| | | 10,827 | 4,942 | 1,227 | 977 |
| | | 4517 | 2924 | 243 | 495 |
| | | 2,117 | 708 | 24 | 246 |
| | | 3,702 | 2,849 | 70 | 147 |
| | | 816 | 279 | 30 | 102 |
| | | 9,701 | 7,422 | 990 | 690 |
| | | 3,646 | 1,370 | 196 | 890 |
| | | 7,436 | 1,426 | 1,235 | 920 |
| | | 3,501 | 1,634 | 158 | 484 |
| | | 6,199 | 2,362 | 2,421 | 1,945 |
| | | 11,833 | 4,317 | 127 | 1,268 |
| | | 13,355 | 7,508 | 591 | 860 |
| | | 5,971 | 1,614 | 114 | 1,151 |
| | | 9,630 | 5,130 | 248 | 1,230 |
| | | 8,360 | 4,706 | 15 | 1,278 |
| | | 2,331 | 1,135 | 152 | 165 |
| | | 14,130 | 12,170 | 123 | 3,742 |
| | | 610 | 312 | 126 | 87 |
| | | 85 | 43 | 49 | 250 |
| | | 1,590 | 6,616 | 350 | 250 |
| | | 3,721 | 1,760 | 485 | 440 |
| | | 3,307 | 1,726 | 56 | 172 |
| | | 4,146 | 2,277 | 53 | 489 |
| | | 5,352 | 3,222 | 187 | 1004 |
| | | 3,203 | 1,946 | 136 | 438 |
| | | 4,836 | 4,179 | 202 | 656 |
| | | 8,894 | 4,097 | 1,265 | 725 |
| 2092 | 2662 | 2662 | 603 | 891 | 1588 |
| 118 | 3935 | 3935 | 1102 | 380 | 532 |
| 533 | 1815 | 1815 | 416 | 33 | 261 |
| 5 | 5093 | 5093 | 1110 | 89 | 630 |
| 39 | 4305 | 4305 | 722 | 8 | 250 |
| 197 | 2809 | 2809 | 873 | 112 | 679 |
| 144 | 5028 | 5028 | 1423 | 126 | 663 |
| 35 | 1371 | 1371 | 411 | 35 | 320 |
| | | 63339 | 33823 | 16 | 367 |
| | | 17990 | 8995 | 0 | 288 |
| | | 14016 | 6271 | 64 | 139 |
| | | 19220 | 20218 | 0 | 185 |
| | | 13973 | 1696 | 132 | 319 |
| | | 8088 | 2504 | 76 | 207 |
| | | 1552 | 747 | 28 | 78 |
| | | 325 | 65 | 320 | 81 |
| | | 24586 | 22305 | 0 | 1081 |
| | | 9684 | 4287 | 0 | 194 |
| | | 21884 | 9529 | 5 | 417 |

| | | | | | |
|---|---|---|---|---|---|
| | | 2989 | 1080 | 98 | 53 |
| | | 29503 | 6341 | | 1128 |
| | | 19443 | 19190 | 0 | 358 |
| | | 30475 | 26057 | 31 | 356 |
| | | 22292 | 10367 | 0 | 530 |
| | | 28483 | 16180 | 9 | 261 |
| | | 16899 | 6000 | 0 | 254 |
