## Supplementary Material 3b for "The value of livestock abortion surveillance in Tanzania: identifying disease priorities and informing interventions"

| Ward | District | Village ID | Cluster 1 (Pastoral) |
| --- | --- | --- | --- |
| Arri | Babati | SEEDZ-17 | 0 |
| Arusha Chini | Moshi Rural | BZ-15 | 0 |
| Engarenaibor | Longido | BZ-13 | 5 |
| Engikaret | Longido | SEEDZ-2 | 22 |
| Kansay | Karatu | SEEDZ-5 | 0 |
| Kikwe | Meru | SEEDZ-14 | 0 |
| Kimokouwa | Longido | SEEDZ-1 | 12 |
| Kindi | Moshi Rural | BZ-03 (BZ-01 - this was a BZ trial ward without HH data); Karanga (BZ-03) next door) | 0 |
| Machame Mashariki | Hai | BZ-12 | 0 |
| Magugu | Babati | SEEDZ-8 | 0 |
| Meserani | Monduli | BZ-16 | 5 |
| Selela | Monduli | BZ-05 | 6 |
| Rau | Moshi Municipality | BZ-06 | 0 |

| Cluster 2 (Agropastoral) | Cluster 3 (Smallholder) | Mode | Village |
|---|---|---|---|
| 6 | 13 | smallholder | Arri |
| 0 | 8 | smallholder | UTAMADUNI |
| 0 | 0 | pastoral | MAIROWA |
| 0 | 0 | pastoral | Engikaret |
| 30 | 0 | agropastoral | Kansay |
| 9 | 10 | smallholder | Nambala |
| 0 | 0 | pastoral | Kimokowa |
| 0 | 8 | smallholder | MAGEREZA |
| 0 | 6 | smallholder | LYAMUNGO SINDE |
| 18 | 2 | agropastoral | Sarame |
| 0 | 0 | pastoral | EMBARARWAI |
| 0 | 0 | pastoral | SELELA |
| 0 | 10 | smallholder | SABASABA |

| Ward | District |
|---|---|
| Arri | Babati |
| Arusha Chini | Moshi Rural |
| Engarenaibor | Longido |
| Engikaret | Longido |
| Kansay | Karatu |
| Kikwe/Nambala village | Meru |
| Kimokouwa | Longido |
| Kindi | Moshi Rural |
| Machame Mashariki | Hai |
| Magugu/Sarame | Babati |
| Meserani | Monduli |
| Selela | Monduli |
| Rau/Uru Kusini | Moshi Municipality |

TOTAL wards where we have livestock data
TOTAL SH wards where we have livestock data
TOTAL AP wardswhere we have livestock data
TOTAL P wards where we have livestock data

**FROM SEEDZ QUESTIONNAIRE**

Abortions in household (over 12 month period)
Abortions in compound (over 12 month period)
Total Abortions
Total Livestock at compounds in SEEDZ HH
Abortions/head of livestock

NOTE: No livestock data from Machame Mashariki through the Ministry census, but figur
NOTE: Rau data is based on average of four Uru wards (former name of Rau) (Takwimu za

| Agroecological | Cattle local | Cattle improv | Cattle total | Goats local | Goats improv | Goats total |
|---|---|---|---|---|---|---|
| SH | | | **5,625** | | | **5,236** |
| SH | 8,125 | | **8,125** | 1,600 | | **1,600** |
| P | 15,889 | | **15,889** | 51,336 | | **51,336** |
| P | 21,798 | | **21,798** | 35,017 | | **35,017** |
| AP | 28,618 | | **28,618** | 14,550 | | **14,550** |
| SH | 21,310 | 4,090 | **25,400** | 19,358 | 2,114 | **21,472** |
| P | 17,585 | | **17,585** | 22,758 | | **22,758** |
| SH | 5,634 | | **5,634** | 1,443 | | **1,443** |
| SH | 6,099 | | **6,099** | | | **5,151** |
| AP | 11,670 | 381 | **12,051** | 11,847 | 51 | **11,898** |
| P | 10,694 | | **10,694** | 10,487 | | **10,487** |
| P | 46,166 | | **46,166** | 42,274 | | **42,274** |
| SH | | | **3,399** | | | **3,313** |
| | | | **207,083** | | | **226,535** |
| | | | **54,282** | | | **38,215** |
| | | | **40,669** | | | **26,448** |
| | | | **112,132** | | | **161,872** |

| Cattle | Goats | Sheep |
|---|---|---|
| 213 | 766 | 418 |
| 337 | 710 | 369 |
| 550 | 1,476 | 787 |
| 22,441 | 12,613 | 10,536 |
| 0.02 | 0.12 | 0.07 |

es for 2023 provided by the LFO (conveyed orally to the research team)

a mifugo Kilimanjaro_v3_combined.xls)

| | | | Reported abortions (SEBI) | | Expected no. abort | |
|---|---|---|---|---|---|---|
| Sheep local | Sheep in | Sheep tota | Cattle | Goats | Sheep | Cattle |
| | | **1,933** | 1 | 1 | 0 | 276 |
| 7,870 | | **7,870** | 11 | 3 | 0 | 398 |
| 17,088 | | **17,088** | 0 | 12 | 12 | 779 |
| 21,135 | | **21,135** | 0 | 25 | 2 | 1068 |
| 4,958 | | **4,958** | 0 | 1 | 0 | 1403 |
| 549 | 192 | **741** | 1 | 1 | 0 | 1245 |
| 15,253 | | **15,253** | 0 | 4 | 1 | 862 |
| 5,987 | | **5,987** | 1 | 0 | 0 | 276 |
| | | **1,433** | 37 | 0 | 0 | 299 |
| | | **4,839** | 0 | 0 | 1 | 591 |
| 7,028 | | **7,028** | 3 | 2 | 0 | 524 |
| 51,908 | | **51,908** | 14 | 42 | 28 | 2263 |
| | | **1,107** | 3 | 9 | 0 | 167 |
| | | | 71 | 100 | 44 | |
| | | **141,280** | 71 | 100 | 44 | 10151 |
| | | **19,071** | 54 | 14 | 0 | 2,006 |
| | | **9,797** | 0 | 1 | 1 | 1,403 |
| | | **112,412** | 17 | 85 | 43 | 5,496 |

ions over 24 month SEBI study % investigated

| Goats | Sheep | Cattle | Goats | Sheep |
|---|---|---|---|---|
| 1225 | 452 | 0.36 | 0.08 | 0.00 |
| 374 | 1842 | 2.76 | 0.80 | 0.00 |
| 12015 | 3999 | 0.00 | 0.10 | 0.30 |
| 8196 | 4947 | 0.00 | 0.31 | 0.04 |
| 3405 | 1160 | 0.00 | 0.03 | 0.00 |
| 5025 | 173 | 0.08 | 0.02 | 0.00 |
| 5326 | 3570 | 0.00 | 0.08 | 0.03 |
| 338 | 1401 | 0.36 | 0.00 | 0.00 |
| 1206 | 335 | 12.38 | 0.00 | 0.00 |
| 2785 | 1133 | 0.00 | 0.00 | 0.09 |
| 2454 | 1645 | 0.57 | 0.08 | 0.00 |
| 9894 | 12149 | 0.62 | 0.42 | 0.23 |
| 775 | 259 | 1.80 | 1.16 | 0.00 |
| 53019 | 33066 | **0.70** | **0.19** | **0.13** |
| 6,703 | 5,423 | **2.69** | **0.21** | **0.00** |
| 3,405 | 1,160 | **0.00** | **0.03** | **0.09** |
| 37,885 | 26,309 | **0.31** | **0.22** | **0.16** |
